# Profiling of open chromatin in developing pig (*Sus scrofa*) muscle to identify regulatory regions

**DOI:** 10.1101/2021.01.21.426812

**Authors:** Mazdak Salavati, Shernae A. Woolley, Yennifer Cortés Araya, Michelle M. Halstead, Claire Stenhouse, Martin Johnsson, Cheryl J. Ashworth, Alan L. Archibald, Francesc X. Donadeu, Musa A. Hassan, Emily L. Clark

## Abstract

There is very little information about how the genome is regulated in domestic pigs (*Sus scrofa*). This lack of knowledge hinders efforts to define and predict the effects of genetic variants in pig breeding programmes. In order to address this knowledge gap, we need to identify regulatory sequences in the pig genome starting with regions of open chromatin. We have optimised the ‘Improved Protocol for the Assay for Transposase-Accessible Chromatin (Omni-ATAC-Seq)’ to profile regions of open chromatin in flash frozen pig muscle tissue samples. This protocol has allowed us to identify putative regulatory regions in semitendinosus muscle from 24 male piglets. We collected samples from the smallest, average, and largest sized male piglets from each litter through five developmental time points. The ATAC-Seq data were mapped to Sscrofa11.1 using Bowtie2 and Genrich was used for post-alignment peak-calling. Of the 4,661 ATAC-Seq peaks identified that represent regions of open chromatin, >50% were within 1 kb of known transcription start sites. Differential read count analysis revealed 377 ATAC-Seq defined genomic regions where chromatin accessibility differed significantly across developmental time points. We found regions of open chromatin associated with down regulation of genes involved in muscle development that were present in small sized foetal piglets but absent in large foetal piglets at day 90 of gestation. The dataset that we have generated provides: i) a resource for studies of genome regulation in pigs, and ii) contributes valuable functional annotation information to filter genetic variants for use in genomic selection in pig breeding programmes.

## Introduction

The domestic pig (*Sus scrofa*) is a hugely important farmed animal species globally, contributing a source of healthy animal protein to feed the growing human population. Meeting the increased demand for healthy sustainably produced food from pigs in coming decades will require novel breeding strategies and management practices that will rely on an improved ability to predict phenotype from genotype (Clark et al. 2020). High resolution annotations of the expressed and regulatory regions of farmed animal genomes provides a resource to accurately link genotype to phenotype (Andersson et al. 2015). Variants in putative regulatory regions have been associated with >100 phenotypes in humans (Pai et al. 2015). Recently, a functional regulatory variant in the gene myosin heavy chain 3 (*MYH3*) was shown to influence muscle fibre type composition in Korean native pigs (Cho et al. 2019). There is very little species-specific information about how the genome is regulated in domestic pigs. This lack of knowledge hinders efforts to identify causative variants for complex traits, and a better knowledge of genome regulation might also improve genomic prediction in breeding programmes. In order to address this knowledge gap, we aim to identify regulatory sequences in the pig genome, starting with regions of open chromatin.

Activation of regulatory DNA drives gene expression patterns that influence the phenotypic characteristics. Measurement of open chromatin gives a quantitative genome wide profile of chromatin accessibility appearing as ‘peaks’ in the data generated for each tissue sample (Thurman et al. 2012). These peaks can reflect the function of the adjoining regulatory DNA (Thurman et al. 2012). The Assay for Transposable Chromatin (ATAC-Seq) (Buenrostro et al. 2013; Buenrostro et al. 2015) has been used successfully to profile regions of open chromatin in chicken, cattle and pig genomes (Halstead et al. 2020a,b). In this study we optimised the ‘Improved Protocol for the Assay for Transposase-Accessible Chromatin (Omni-ATAC-Seq)’ (Corces et al. 2017) to profile regions of open chromatin in flash frozen pig muscle tissue samples.

Muscle is an important tissue in commercial pig production as muscle traits (e.g. meat and carcass quality) act as economic drivers in pig breeding programmes. Prior to this study knowledge of open chromatin in pig muscle was limited to data from only two adult animals (Halstead et al. 2020a) and four foetuses from three early developmental stages (Yue et al. 2021). For this study, we collected semitendinosus muscle tissues from piglets at five different stages of development (three foetal stages, one neonatal and one juvenile stage). The developmental stages were chosen according to their relevance to hyperplasic muscle development in the foetus and post-natal muscle hypertrophy (Ashmore et al. 1973; Wigmore and Stickland 1983; Rudar et al. 2019). We hypothesised that gene expression and regulation in semitendinosus muscle tissue would change as the piglets aged, allowing us to identify the transcripts and regions of open chromatin that drive myogenesis. Several studies have profiled gene expression during foetal development in pigs (Zhao et al. 2011; Yang et al. 2015; Zhao et al. 2015; Ayuso et al. 2016), however to date only one other study has examined how chromatin openness changes as the piglet develops (Yue et al. 2021).

The number of muscle fibres in pigs is proportional to weight at birth (Aiello et al. 2018; Stange et al. 2020). Low birth weight in pigs has been shown to cause lifelong impairments in muscle development and growth (Rehfeldt and Kuhn 2006). Low birth weight piglets often display ‘catch up’ growth, but at the expense of laying down a higher proportion of body fat compared to normal sized littermates (Estany et al. 2017). Consistent with these observations, mesenchymal stem cells from intrauterine growth-restricted piglets show a differentiation bias towards the adipocyte lineage in comparison with their normal sized litter mates (Weatherall et al. 2020). Low birth weight piglets tend to produce fatter, less valuable carcasses from a production perspective and as such their incidence within pig litters should be kept to a minimum (Pardo et al. 2013). Piglet size variation within a litter is likely to be determined by many different physiological variables including variation in placental blood flow (Stenhouse et al. 2018) but may also be influenced by genetic and epigenetic factors (Wang et al. 2016; Li et al. 2020).

The study we present here used samples of muscle tissue from a common commercial breed cross (Large White x Landrace) to generate ATAC-Seq and RNA-Seq data from the same individuals to characterise the expressed and regulatory regions of the genome during pig development. The aims of the study were to: 1) Optimise the Omni-ATAC-Seq protocol for frozen pig muscle tissue; 2) Map regions of open chromatin in semitendinosus muscle tissue from small, average and large sized male piglets at five developmental stages (days 45, 60 & 90 pre-natal, one and six weeks post-natal) and 3) Analyse RNA-Seq data from the same tissues to generate gene expression profiles. To our knowledge, this is the first time the Omni-ATAC-Seq protocol has been optimised for frozen muscle tissue from a farmed animal species and we have provided a detailed protocol on the Functional Annotation of Animal Genomes (FAANG) data portal (https://data.faang.org). The outcomes of the study will help to 1) Understand the molecular drivers of muscle growth in pigs; 2) Provide a foundation for functionally validating target genomic regions *in vitro* and 3) Identify high quality causative variants for muscle growth with the goal of harnessing genetic variation and turning it into sustainable genetic gain in pig breeding programmes.

## Methods

### Animals

Tissue samples for this study were collected from Large White x Landrace pigs that were euthanized, not specifically for this study but for other on-going projects on the effects of foetal size on pig development at The Roslin Institute. Foetal tissues were collected from pregnant sows that were euthanized with sodium pentobarbitone 20% w/v (Henry Schein Animal Health, Dumfries, UK) at a dose of 0.4 ml/kg by intravenous injection. Post-natal samples were collected after euthanasia by captive bolt.

### Sample collection of frozen muscle tissue samples for ATAC-Seq and RNA-Seq

The tissue samples used for this study were from archived material (with the exception of the piglets that were six weeks of age) collected from the largest, smallest and average sized male piglets per litter at five different developmental stages (Table 1). The largest, smallest and average sized piglets from each litter were selected according to body weight for the foetal time points and birth weight for post-natal time points (Supplementary Table 1). Developmental stages were chosen, according to previous studies (Ashmore et al. 1973; Wigmore and Stickland 1983; Rudar et al. 2019) as follows:

**Table 1.**
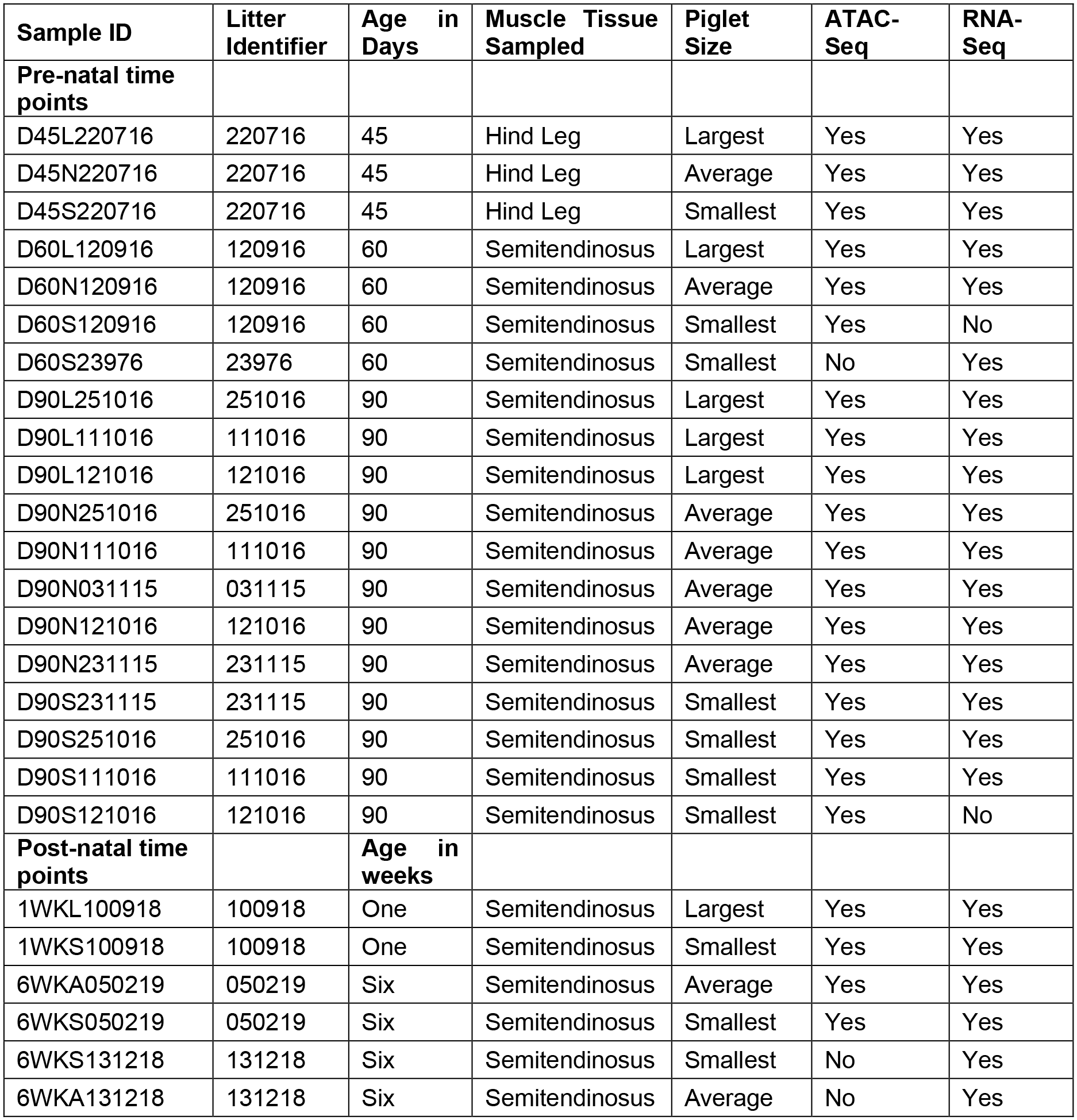
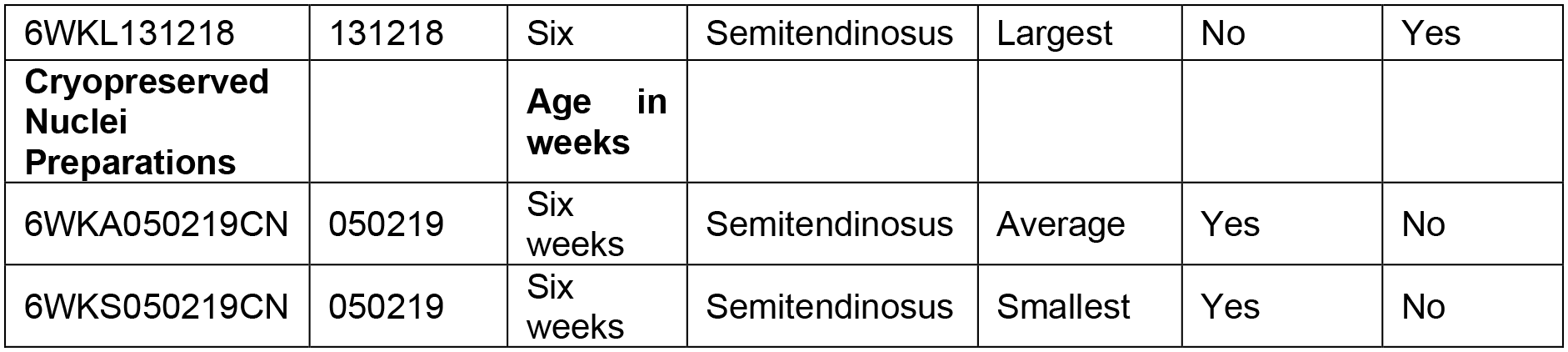
Details of muscle tissues sampled, for ATAC-Seq and RNA-Seq, from piglets at five developmental stages.

Day 45 of gestation - when primary muscle fibres form.

Day 60 of gestation - when secondary muscle fibres begin to form.

Day 90 of gestation - when fibre formation ceases after which subsequent muscle growth occurs through fibre hypertrophy.

One week of age - during active muscle hypertrophy.

Six weeks of age - once muscle hypertrophy has levelled off.

Due to limited sample availability the experimental design is unbalanced (Table 1). Only one complete set of littermates (smallest, average and largest) were included in the analysis for the day 45 (n=3) and 60 (n=3) time points. For the six weeks of age time point (n=5) one large sample was unavailable for inclusion in the analysis. At day 90 (n=11) of gestation three complete litters and one incomplete litter were included, while at one week of age samples were only available from the smallest and largest piglets from one litter (n=2). We have included a flow chart describing which samples were analysed at each stage of this study (Figure 1).

**Figure 1:**
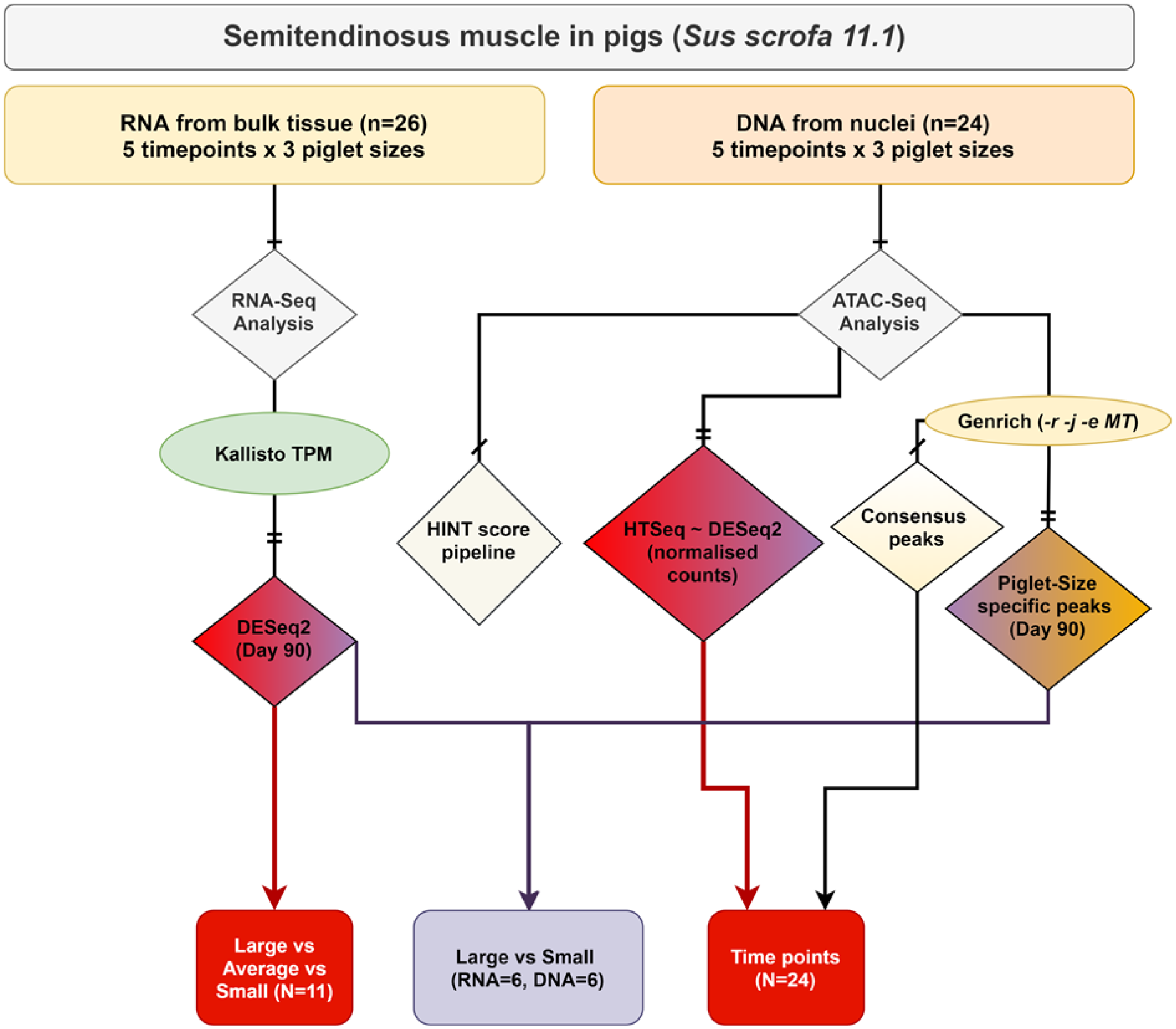
Flowchart describing the experimental design and samples included in each stage of the analysis performed in this study. Colour coding indicates where there are overlaps in the analysis performed for each component of the study.

Samples were collected from the semitendinosus muscle from the hind leg of piglets from each developmental stage (Table 1). The only exception was day 45, when whole muscle tissue was collected, because it was not possible to differentiate specific muscle types at this early stage of development. Each sample was flash frozen in liquid nitrogen, as quickly as possible within an hour post euthanasia and stored at −80°C for future analysis. From the six week old piglets additional samples were collected in sucrose buffer to isolate and cryopreserve nuclei according to the method described in (Halstead et al. 2020a). Cryopreserving isolated nuclei for a small number of samples would allow us to validate the data we generated from the flash frozen material, which we were optimising for the first time on muscle tissue. The protocol for collection of tissue samples at the farm is available via the FAANG Data Coordination Centre https://data.faang.org/api/fire_api/samples/ROSLIN_SOP_Collection_of_tissue_samples_for_ATAC-Seq_and_RNA-Seq_from_large_animals_20200618.pdf.

### Isolation of cryopreserved nuclei from fresh muscle tissue and preparation of tagmented nuclear DNA

We used the protocol described in (Halstead et al. 2020a) to isolate and cryopreserve intact nuclei from fresh muscle tissue samples from the six week old piglets (Table 1). Briefly, each tissue sample was transferred to a GentleMACS C tube (Mitenyi Biotec, Germany) with sucrose buffer and homogenised. The homogenate was then filtered and Dimethyl Sulfoxide (DMSO) (Sigma Aldrich, USA) added (10% final concentration), before freezing at −80°C overnight in a Mr Frosty (Nalgene, USA), then transferring to a −80°C freezer for long-term storage. The full protocol for preparation of cryopreserved nuclei from fresh muscle tissue is available via the FAANG Data Coordination Centre https://data.faang.org/api/fire_api/samples/ROSLIN_SOP_Cryopreservation_of_Nuclei_for_ATACSeq_using_GentleMACS_20201119.pdf.

To prepare tagmented DNA the cryopreserved nuclei preparations were thawed slowly at room temperature by adding 500 µl of cold 1x Phosphate Buffered Saline (PBS), filtered then centrifuged at 500 x g at 4°C in a swinging bucket centrifuge for 10 minutes. After centrifugation, the pellet was resuspended in 1 ml cold ATAC-Seq RSB buffer + 0.1% Tween20 (Sigma Aldrich, USA) for lysis and centrifuged for 10 minutes at 500 x g at 4°C. The pellet of nuclei was then washed in PBS and resuspended in 50 µl transposition mix (25 µl TD buffer, 2.5 µl TDE1 enzyme, Molecular Biology Grade Sterile H_2_O) from the Nextera DNA Sample Prep Kit (Ilumina, USA). The pellet was incubated with the transposition mix for 60 minutes at 37°C at 300 rpm on a thermomixer. The pellet of transposed nuclear DNA, was purified with a MinElute PCR purification kit (Qiagen, Germany), eluted in 15 µl of Buffer EB, and stored at *-*20°C. The full protocol is available via the FAANG Data Coordination Centre https://data.faang.org/api/fire_api/samples/ROSLIN_SOP_ATAC-Seq_DNAIsolationandTagmentation_Cryopreserved_Muscle_Nuclei_Preparations_20200720.pdf

### Isolation of nuclei from frozen muscle tissue and preparation of tagmented nuclear DNA

ATAC-Seq libraries were prepared using a version of the ‘Improved Protocol for the Assay for Transposase-Accessible Chromatin (Omni-ATAC-Seq)’ (Corces et al. 2017) which we optimised for flash frozen pig muscle tissue samples for this study. The main modification that we introduced to the protocol was an initial dissociation step using a GentleMACS Dissociator (Mitenyi Biotec, Germany), essentially combining the Omni-ATAC-Seq protocol with the initial steps from (Halstead et al. 2020a). The protocol is described in full at https://data.faang.org/api/fire_api/samples/ROSLIN_SOP_ATAC_Seq_DNAIsolationandTagmentation_Frozen_Muscle_Tissue_20200720.pdf and summarised here. The components of each of the buffers are included in Supplementary Table 2. Each flash frozen tissue sample (∼200 mg per sample) was chopped into small pieces over dry ice and then dissociated in a GentleMACS C-tube (Mitenyi Biotec, Germany) in 1 ml of 1XHB buffer (+Protease Inhibitor Cocktail (PIC)). The samples were dissociated using programme m_muscle_0.1_0.1 (equivalent to ‘E0.1c Tube’) twice on a GentleMACS Dissociator (Mitenyi Biotec, Germany). Immediately after dissociation the samples were filtered through a 70 μm corning cell strainer (Sigma Aldrich, USA) then centrifuged at 3000 x g for 5 minutes. The pellet was resuspended in 400 μl 1XHB buffer and transferred to a 2 ml Eppendorf Protein Lo-Bind tube (Eppendorf, UK). 400 μl of 50% Iodixanol solution (Opti-Prep Density Gradient Medium) (SLS, UK) was added to the 400 μl of cell solution (final 25% Iodixanol). An Iodixanol gradient was then created and samples transferred to a swinging bucket centrifuge and spun for 25 minutes at maximum speed at 4°C with no brake. A thin “whitish” band appeared between layers two and three of the gradient. Evaluation and counting of nuclei was performed by staining with Trypan Blue (ThermoFisher Scientific, USA). 1 ml of ATAC-RSB Buffer + 0.1% Tween20 (Sigma Aldrich, USA) was then added to lyse the nuclei and the sample centrifuged for 10 minutes at 500 x g at 4°C. The pellet was then gently resuspended in 50 µl transposition mix for tagmentation as described for cryopreserved nuclei samples above.

### ATAC-Seq library preparation

The library preparation protocol, adapted from (Corces et al. 2017), was used for the flash frozen tissues and the cryopreserved nuclei preparations. The protocol described in full is available via the FAANG Data Coordination Centre https://data.faang.org/api/fire_api/samples/ROSLIN_SOP_ATAC-Seq_LibraryPreparationandSizeSelection_20200720.pdf. A PCR reaction mix was set up comprising 10 μl molecular biology grade H_2_O, 2.5 µl Ad1 primer 25 µM, 2.5 µl Ad2.x primer 25 µM (variable index see Supplementary Table 3), and 25 µl 2x NEBNext Hi-Fi PCR mix (NEB, USA) per reaction. 10 µl of transposed DNA was added to each reaction and 5 amplification cycles of the following PCR reaction performed: 72°C for 5 min, 98°C for 30 sec, 98°C for 10 sec, 63°C for 30 sec, 72°C for 1 min. The GreenLeaf Quantitative PCR (qPCR) Protocol (Buenrostro et al., 2015) was used to determine the number of additional PCR amplification cycles that were required for each sample, to stop amplification prior to saturation and avoid variation across samples caused by PCR bias. Samples for which more than 5-7 additional cycles were required were discarded due to the high probability of PCR bias caused by additional cycles. Amplified ATAC-Seq libraries were then purified with a MinElute PCR purification kit (Qiagen, Germany). Library quality was checked on the Agilent 2200 TapeStation System (Agilent Genomics, USA). Libraries were assessed for quality according to an even distribution of fragments and a clearly differentiated sub-nucleosomal fragment as described in (Halstead et al. 2020a). If library quality was sufficient the sub-nucleosomal fragment (150-250 bp) was size selected, in order to minimise the signal to noise ratio, as suggested in (Halstead et al. 2020a). Size selection was performed using a Thermo Scientific E-Gel System (ThermoFisher Scientific, USA). To check the size of the selected fragment an aliquot was run on the Agilent 2200 TapeStation System (Agilent Genomics, USA). After size selection the libraries were pooled and stored at −20°C prior to sequencing.

### Sequencing of ATAC-Seq libraries

Pooled libraries (4 batches) were sequenced to generate 50 nt paired-end reads on an Illumina NovaSeq 6000 platform using a single S2 flow cell. All of the libraries generated >90M paired-end reads (Min: 9.8e+07, Max: 3.5e+08, Median: 1.97e+08).

### ATAC-Seq data processing and mapping

Quality control of the raw sequence data was performed using FastQC v0.11.9 (Andrews 2010) and multiQC v1.9 (Ewels et al. 2016). The paired end reads were trimmed using Trimmomatic v0.39 (Bolger et al. 2014). The trimmed reads were then mapped to the Sscrofa11.1 pig reference genome (Warr et al. 2020) available from Ensembl (GCA_000003025.6) using Bowtie2 v2.3.5.1 and the default flags of the *--very-sensitive* mode followed by excluding unmapped reads and marking PCR duplicates. PCR duplicates were marked using Picardtools v2.23.0 (Li et al. 2009; Broad Institute 2019). The BAM files that were generated were then sorted and indexed using samtools v1.6 (Li 2011). Overall on average more than 75 M reads per samples were uniquely mapped (Min: 2.47e+07, Max: 1.28e+08, Median: 7.74e+07, Mean ± SD: 7.72e+07 ± 3.15e+07). The PCR duplication level (post mapping) was 43% ± 8 (mean ± SD) across all libraries.

The following parameters were measured as recommended by the ENCODE project for validation of ATAC-Seq libraries (Davie et al. 2015; Davis et al. 2018): Fragment size distribution (PicardTools v2.25.4 (Broad Institute 2019)), Fragment in peak (FRiP score, deeptools v3.5.1 (Ramírez et al. 2016)), transcription start site enrichment score (TSS-ES ATACseqQC v1.14.4 (Ou et al. 2018)), Non-redundant fraction (NRF samtools v1.6 (Li et al. 2009)), nucleosome free region score (NFR score ATACseqQC v1.14.4 (Ou et al. 2018)) and PCR bottlenecking coefficient (PBC ATACseqQC v1.14.4 (Ou et al. 2018)). A detailed table of QC scores is included in Supplementary Table 6.

### ATAC-Seq peak calling using Genrich

ATAC-Seq peak calling for each developmental time point was performed using Genrich v0.5 (John M. Gaspar 2020) under ATAC-Seq mode while excluding PCR duplicates and mitochondrial reads (used flags: *-r-j-e MT*). Two rounds of peak calling were performed as follows: 1) peak calling on individual samples (n=24), 2) aggregated multi-sample peak calling for each piglet size. All the scripts used for this analysis can be found in Supplementary File 1 and the code repository https://msalavat@bitbucket.org/msalavat/pig_muscle.git.

### Multidimensional scaling analysis for comparison of ATAC-Seq libraries

Non-linear multidimensional scaling (NMDS) of the ATAC-Seq libraries was performed using the MASS::IsoMDS package (Venables and Ripley 2002) to ensure that there were no obvious outlying samples and that tissues of the same type clustered together in a biologically meaningful manner. A distance matrix (Manhattan distance) was produced using the consensus set of ATAC-Seq peaks and the normalized read counts as described in the previous section. The distance matrix was then processed for multidimensional scaling. NMDS was also used to compare the ATAC-Seq libraries prepared from either flash frozen muscle tissue or cryopreserved nuclei from two 6-week-old piglets.

### Differential peak analysis based on read counts

A consensus set of ATAC-Seq peaks was created for the purpose of differential peak analysis. The consensus set, from individual sets that were called in all 24 samples, was created using bedtools v2.26.0 (*bedtools merge -i all_samples.bed -d10 -c 4,7,10,4 -o count_distinct,mean,mode,distinct*). Peaks that were within <=10 nucleotides of another peak were merged in to one peak, and a support value (i.e., the number of tissue samples in which the peak was present) was calculated for each peak. Peaks with a support value of less than 3 (i.e., they were present in less than 3 tissue samples) were removed, resulting in a total of 12,090 ATAC-Seq peaks, which will now be referred to as the “consensus set”. A read fragment filtering and analysis workflow was devised similar to a recently published framework by (Yan et al. 2020). Briefly, the mapped BAM files were filtered for high mapping quality, non-PCR duplicates and non-mitochondrial reads using samtools v1.6 (*samtools view -h -f2 -q10 -F1548 -bS*). A read count for each sample (using high quality BAM files) was then generated using ht-seq v0.13.5 (Anders et al. 2015) against the consensus set of ATAC-Seq peaks (*htseq-count --stranded=no –type=region*). The library size for each BAM file was then used to normalise the read counts for multidimensional scaling analysis as described by (Yan et al. 2020) using the following equation:

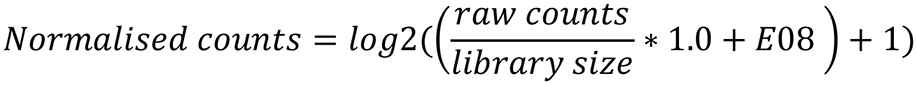

DESeq2 v1.30.1 (Love et al. 2014) was used for differential peak analysis, of the raw read counts, to compare across developmental time points. A Likelihood Ratio Test (reduced model) was used for the analysis of the time points (design: ∼ size + time; reduced: ∼ size). A multiple testing *p* value correction was performed using the Benjamini-Hochberg (Benjamini and Hochberg 1995) method and a 10% false discovery rate (FDR) was considered as the threshold of significance.

### Transcription factor footprint analysis

The HMM-based Identification of Transcription factor footprints (HINT) pipeline from the Regulatory Genomics Toolbox (RGT; v0.12.3) (Li et al. 2019) was used to compare transcription factor (TF) activity between developmental stages or piglet sizes. For a given comparison, the rgt-hint command in footprinting mode was used to identify TF footprints within peaks based on ATAC-Seq signal in each condition. When comparing consecutive developmental stages, the ATAC-Seq peaks identified for each stage were merged with Bedtools (v2.26.0), and footprints were identified within the merged set. When comparing different piglet sizes within a developmental time point, footprints were identified within the peak set for that time point (regardless of piglet size). ATAC-Seq signal for a given condition included aligned reads from all biological replicates (excluding libraries from cryopreserved nuclei), which were combined and filtered to remove duplicates using Samtools (v1.7). Footprints were matched to known motifs in JASPAR (Fornes et al. 2020) with rgt-motif analysis, and rgt-hint in differential mode was then used to compare the activity of each TF between two given conditions using bias-corrected signal.

### RNA isolation and quality control

The RNA isolation protocol is described in full at https://data.faang.org/api/fire_api/samples/ROSLIN_SOP_RNA_IsolationoftotalRNAfromfrozentissuesamples_20200720.pdf. RNA was extracted from approximately 60mg of tissue. Tissue samples were homogenised in 1 ml of TRIzol (Thermo Fisher Scientific, USA) with CK14 (VWR, USA) tissue homogenising ceramic beads on a Precellys Tissue Homogeniser (Bertin Instruments; France) at 5000 rpm for 20 sec. RNA was then isolated using the TRIzol protocol (Thermo Fisher Scientific, USA) and column purified to remove DNA and trace phenol using a RNeasy Mini Kit (Qiagen, Germany) following the manufacturer’s instructions. RNA integrity (RIN^e^) was estimated on an Agilent 2200 TapeStation System (Agilent, USA) to ensure RNA quality was of RIN^e^ > 7. RIN^e^ and other quality control metrics for the RNA samples are included in Supplementary Table 4.

### Poly-A enriched library preparation and sequencing

Strand-specific paired-end reads with a fragment length of 100 bp for each sample were generated by Edinburgh Genomics, using the Illumina TruSeq mRNA library preparation protocol (poly-A selected) (Illumina; Part: 15031047 Revision E). mRNA-Seq libraries were sequenced on an Illumina NovaSeq 6000 platform to generate >66 M paired end reads per sample (Min: 6.6e+07, Max:1.21e+08, Mean: 9.17+e07).

### RNA-Seq data analysis workflow

The raw sequence data were quality controlled and trimmed using Trimmomatic (Bolger et al. 2014). The Kallisto aligner (Bray et al. 2016) was used for expression quantification of the RNA-Seq data. Briefly, a reference transcriptome fasta file of coding sequences was obtained from Sscrofa11.1 Ensembl v100 to build a Kallisto index file using default settings. The trimmed reads were then mapped for transcript level expression quantification (*de novo)* in kallisto with *--bias* option activated. The output tab separated value files were then imported to R using txImport package (Soneson et al. 2016) for further analysis and visualisations.

The TPM expression estimates for each sample were investigated using principal component analysis (PCA) in FactoMineR to identify any spurious samples that did not cluster as expected (Lê et al. 2008). Differential expression (DE) analysis was performed only on the three sizes of foetal piglet (small, average and large) at day 90 of gestation. The Likelihood Ratio Test (LRT) model of DESeq2, including post-hoc analysis, was used with small size as the reference level i.e. denominator in log2FC (*DESeqDataSetFromTximport(txi = dds,design = ∼ Piglet size)*). After multiple correction of *p* values using the BH method (Benjamini and Hochberg 1995), a false discovery rate of 10% was considered as the significance threshold.

### Overlay of differentially expressed genes and ATAC-Seq peaks

For the day 90 samples only, we re-analysed the ATAC-Seq peaks in the three sizes of foetal piglet per litter (large, average and small). Peak calling was performed using the same Genrich flags as previously described (aggregated multi-sample method), and we separated peaks shared between all size classes and size class specific peaks with bedtools v2.26.0 (Quinlan and Hall 2010). The size specific peaks generated for the foetal piglets at day 90 of gestation and the scripts used to produce them can also be found in Supplementary File 1 and the code repository https://msalavat@bitbucket.org/msalavat/pig_muscle.git.

An overlay of genes that were differentially expressed between the large and small sized foetal piglets at day 90 was performed using ATAC-Seq peaks within 10 kb vicinity of the differentially expressed genes (either upstream or downstream). This overlay would show us which of the differentially expressed genes had an ATAC-Seq peak in their vicinity and whether that peak was present in both large and small sized foetal piglets, or only in one of the two sizes. The distance from the start of the gene model to the start of the ATAC-Seq peak was used as a coordinate system (i.e. positive values meant the peak was either within the gene or within the 5’ 10 kb upstream region of the gene, and negative values corresponded to 10 kb from the 3’ end of the gene).

### Statistical analysis software and packages

All data analysis for this study was performed via bash scripting and use of R (R Core Team 2017) on the University of Edinburgh research computing facility (Edinburgh 2020). The data analysis protocol for ATAC-Seq and RNA-Seq are available at https://data.faang.org/api/fire_api/analyses/ROSLIN_SOP_ATAC-Seq_analysis_pipeline_20201113.pdf and https://data.faang.org/api/fire_api/analyses/ROSLIN_SOP_RNA-Seq_analysis_pipeline_20201113.pdf.

## Results

### ATAC-Seq data from frozen pig muscle tissues

ATAC-Seq libraries from four different batches (24 samples in total) were multiplexed and sequenced to achieve 2.02e+08 average reads per sample (Min: 9.8e+07, Max: 3.5e+08, Median: 1.97e+08). Reads were evenly distributed between barcodes across the first three batches. The fourth batch, which included only two samples that had a higher concentration of starting DNA, resulted in more reads per sequencing run compared to the other 22 samples (details in Supplementary File 2). Average chromosomal coverage across autosomes was 7.2x, 4x for X,3.2x for Y and 4.49e+04 for the mitochondrial chromosome (Supplementary Figure S1). Visual comparison of the ATAC-Seq and the RNA-Seq reads, mapped to the Sscrofa11.1 genome, was used to check for consistency between the two datasets. For example, Figure 2 shows the ATAC-Seq and RNA-Seq data as parallel tracks for a housekeeping gene (*GAPDH* Figure 2A) and a gene related to muscle development (*CASQ1* Figure 2B). Read coverage of all 24 ATAC-Seq samples and the consensus peak called for *GAPDH* has also been visualised in Supplementary Figure S2.

**Figure 2:**
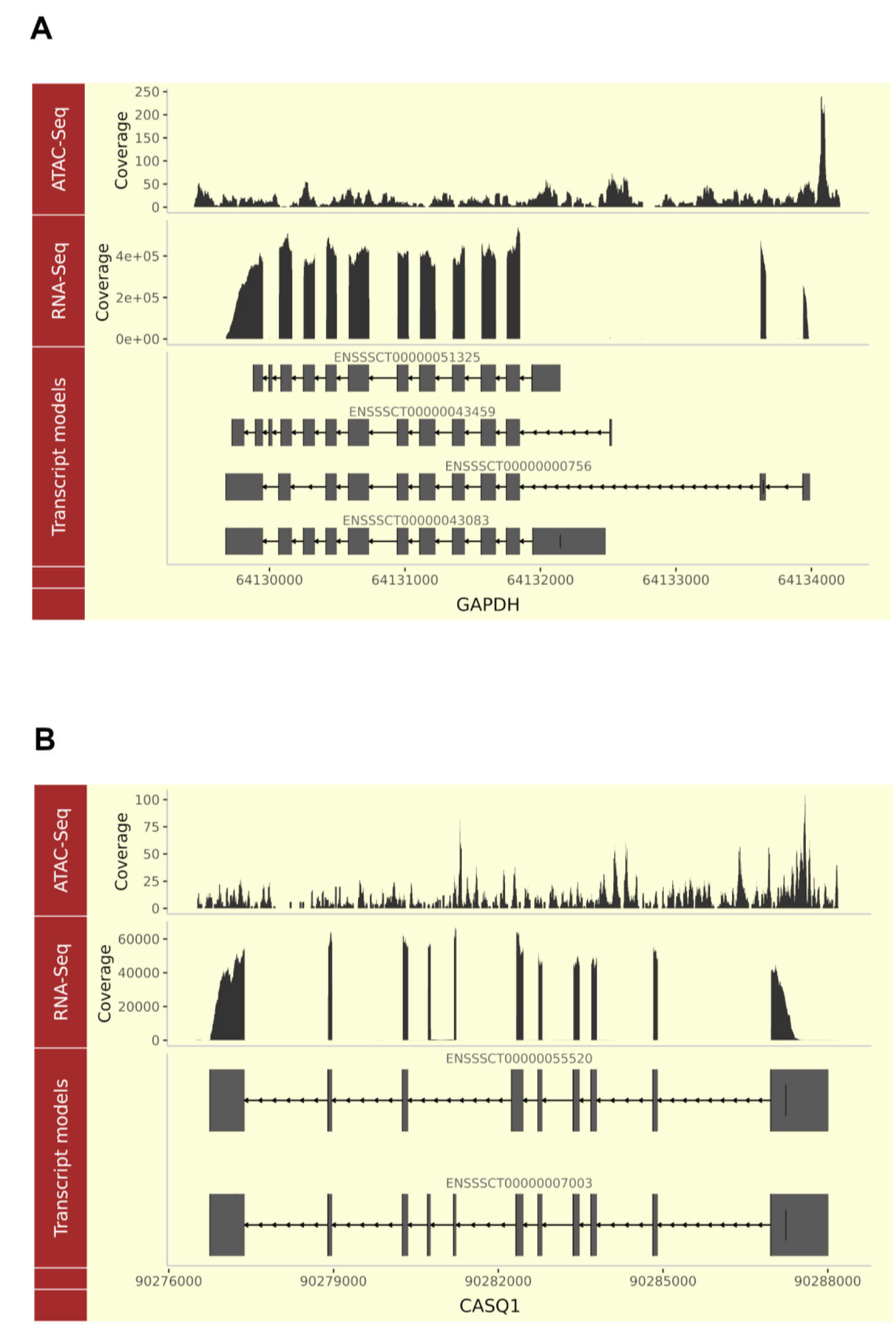
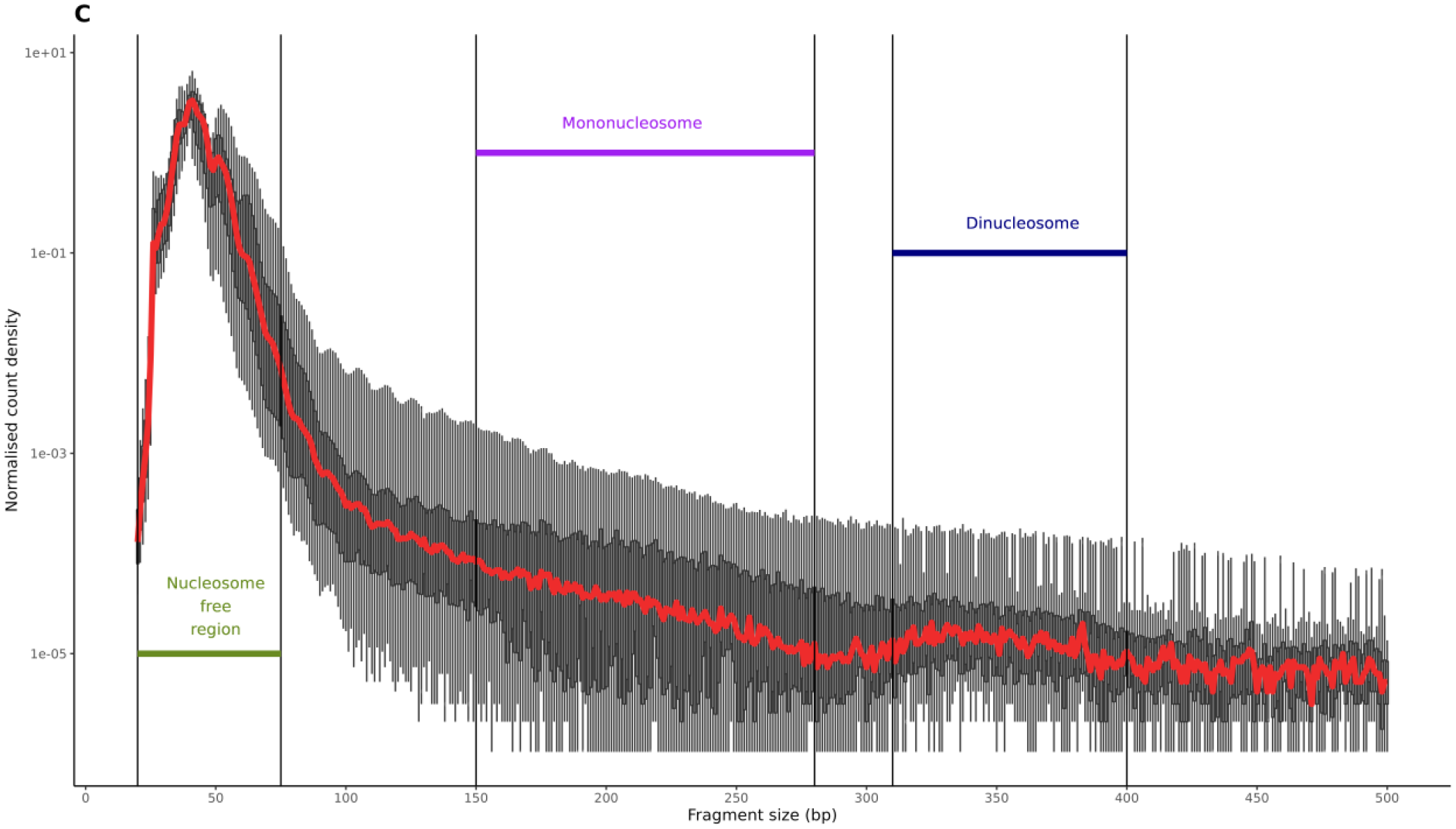
Genomic track visualisation of the ATAC-Seq and RNA-Seq datasets by presence of the signal at the genomic coordinates of two genes GAPDH and CASQ1. The raw ATAC-Seq read counts and RNA-Seq TPM counts, from a representative with a library size closest to the average for the set, are shown in tracks above each gene’s transcript models. Two genes are shown: A) GAPDH a house-keeping gene and B) CASQ1 a gene involved in muscle growth and development. Two additional tracks from a house-keeping gene and gene involved in muscle growth and development and included in Supplementary Figure S2. C) The fragment size distribution of all 24 libraries are plotted against normalised read count density (log2(count/max(count)). Fragment size distribution was calculated after removing PCR duplicates, multi-mapped or improper pairs or mapping quality <30.

We also measured the fragment size distribution of the 24 ATAC-Seq samples, which showed read density was highest at the putative nucleosome free region (approximately 50bp insert size) followed by the mononucleosome region (∼150-200bp) and then the dinucleosome (∼300-400bp) region as shown in Figure 2C.

### Multidimensional scaling analysis of the ATAC-Seq libraries from frozen tissue samples

Non-linear multidimensional scaling (NMDS) was used to ensure that the ATAC-Seq dataset was biologically meaningful, reproducible and there were no outlying samples (i.e. samples from the same developmental stage should have a similar peak distribution and cluster together). NMDS was performed using the consensus set of peaks and the normalised ATAC-Seq read counts. The input matrix, which was converted to a Manhattan distance matrix prior to analysis, consisted of 12,090 consensus peaks and 24 samples. The sample separation on the first two components of the NMDS was driven by the developmental time points. The second and third components of the plot showed the greatest segregation between the developmental time points as shown in Figure 3. A correlation heatmap of the 24 ATAC-Seq samples also indicated that the samples for each time point clustered closely together (Figure 3C). The heatmap input matrix was based on the deeptools v3.5.1 multiBamSummary binned (100bp) read coverage.

**Figure 3:**
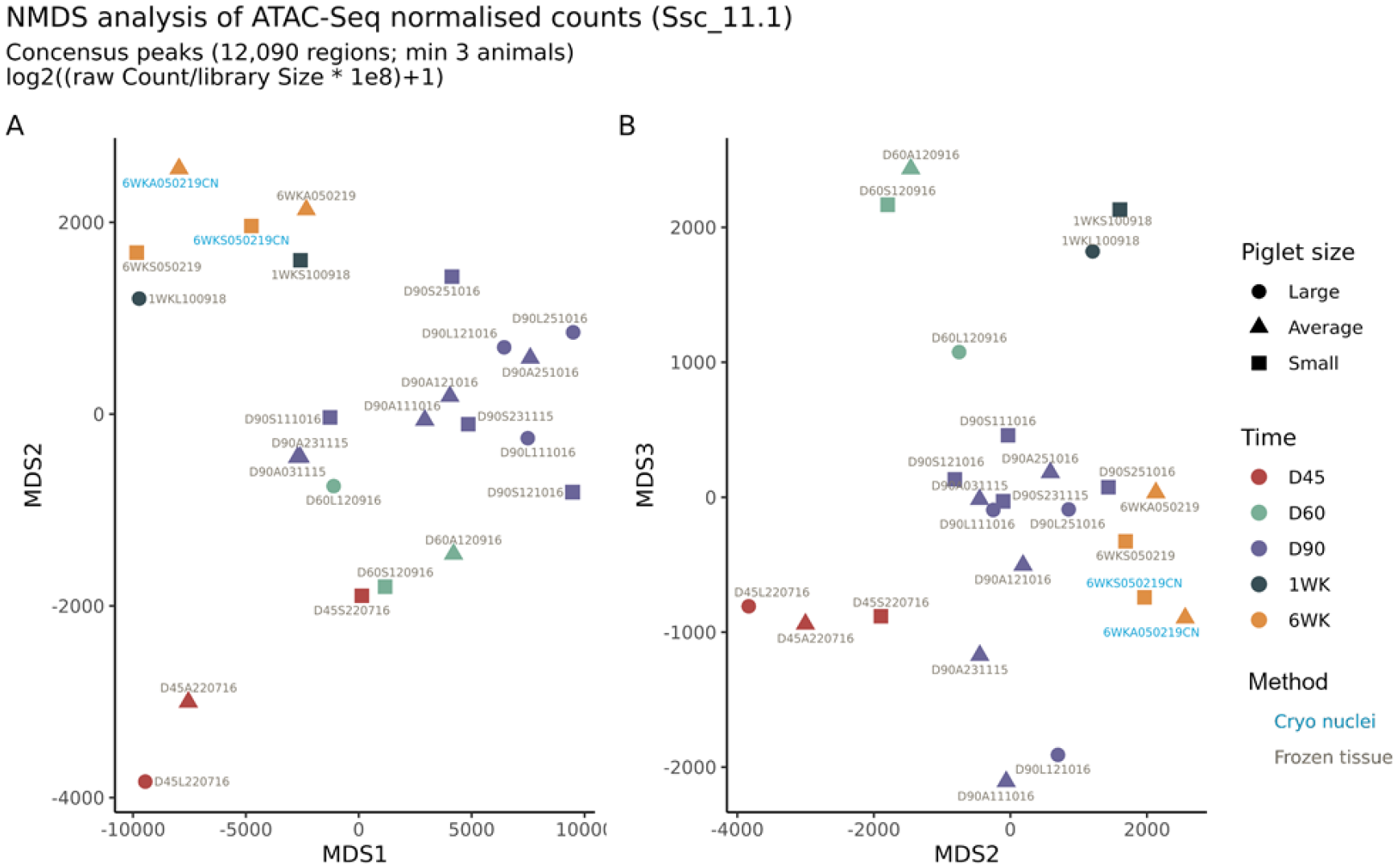
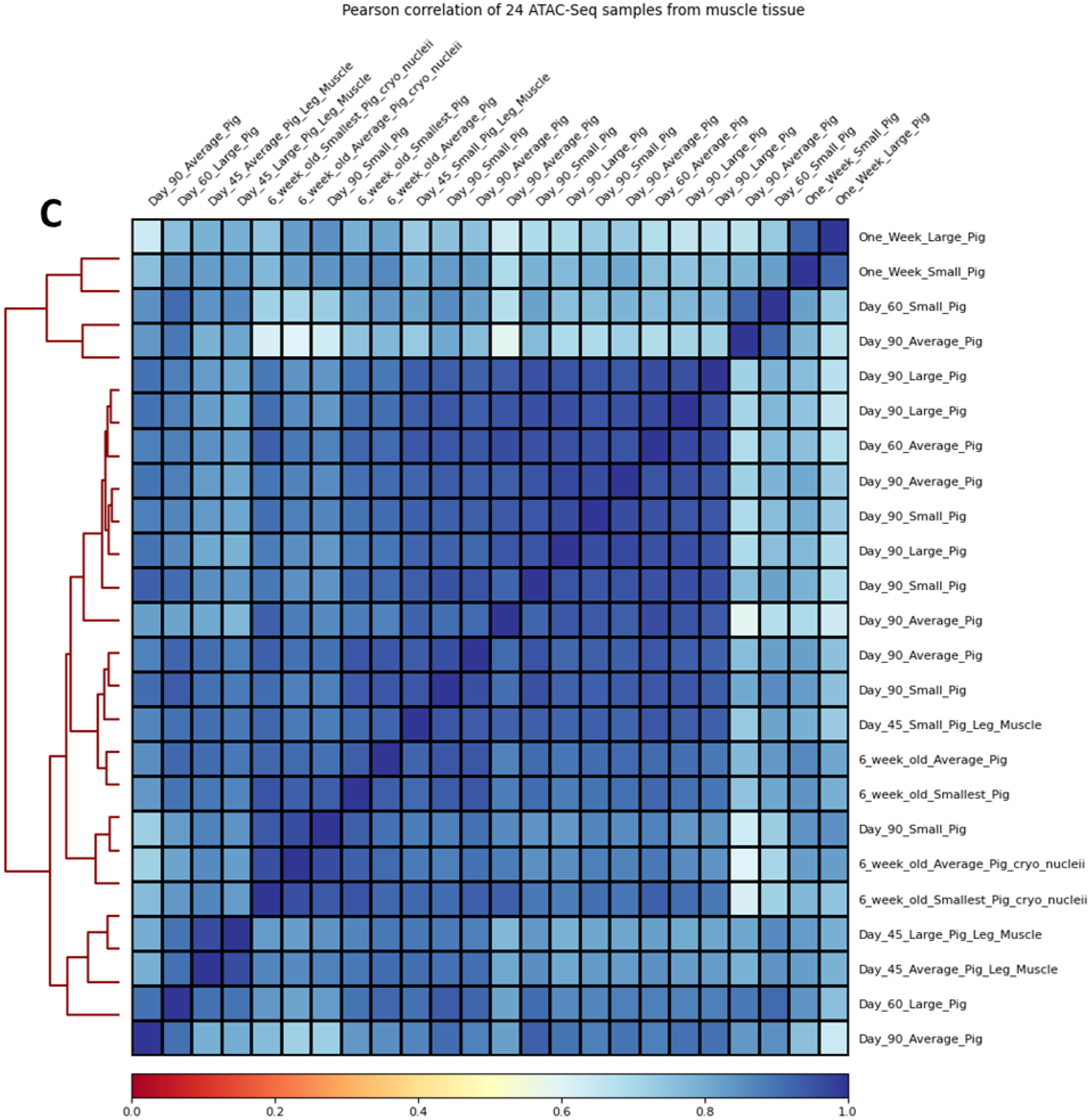
Non-linear Multi-dimensional scaling (NMDS) and correlation analysis of the ATAC-Seq open chromatin consensus set in all samples. A & B) The normalised read count for each was used as described by (Yan et al. 2020).Samples from different developmental time points are indicated by colour and piglet size by shape. Label colours are used to differentiate between Cryopreserved nuclei and Frozen tissue library preparation protocols. C) Correlation heatmap of all 24 samples based on deepTools v3.5.1 binned read coverage (multiBamSummary).

### Multidimensional scaling analysis of ATAC-Seq libraries prepared from either flash frozen muscle tissue or cryopreserved nuclei

Non-linear multidimensional scaling (NMDS) was also used to compare ATAC-Seq libraries prepared from either flash frozen muscle tissue or cryopreserved nuclei from two 6-week-old piglets. The two libraries prepared for the cryo-preserved nuclei samples clustered closely with the libraries prepared for flash frozen tissue indicating there was little difference in the data generated by the two protocols (Figure 3). Other metrics, including the percentage of ATAC-Seq peaks within promoter, proximal, distal regions or within a gene model were also used to compare the libraries prepared from cryopreserved nuclei and flash frozen tissue (Supplementary Figure S3). For each of the metrics chosen libraries prepared from cryopreserved nuclei and flash frozen tissue appeared broadly similar, with a slightly higher percentage of ATAC-Seq peaks in promoter regions (1<=1 kb) in cryopreserved nuclei and in distal intergenic regions in the libraries prepared from frozen tissues (t.test; *p* = 0.4) (Supplementary Figure S3). There were no statistically significant differences detected between the two protocols for any of the ATAC-Seq quality control metrics chosen (ANOVA; *p* > 0.05) except the TSS enrichment score (TSS-ES) (t.test *p* = 0.045; cryo nuclei vs frozen tissue mean TSS-ES 8.5 vs 5.97; Supplementary Table 6 and Supplementary Figure S4).

### Distribution of ATAC-Seq peaks within genomic features

The feature distribution of the ATAC-Seq peaks in all 24 samples is shown in Figure 4A. On average > 6,500 peaks (including overlapping regions) were called in each sample (Min: 2.34e+03, Max: 1.42e+04,Median: 6.38e+03, Mean ± SD: 6.72e+03 ± 3.63e+03). More than 52% of the peaks were located in promoter regions in the majority of samples (19/24). There was a slight negative trend between increase in library size (depth of sequencing) and in the number of the peaks called (linear regression: slope= −6e-05 and R² = 0.22). Detailed metrics can be found in Supplementary File 2. In five samples (day 90 Large [n=3], a day 90 Average [n=1] and a day 90 Small [n=1]) the majority of peaks were located in distal intergenic regions (Figure 4A). We could not find any batch effect, in nuclei extraction or library preparation that might account for this and as such concluded that this variation was related to the samples themselves. There was also little observable difference in how the ATAC-Seq peaks were distributed within the genome of the libraries from cryopreserved nuclei relative to the libraries from flash frozen tissue (Figure 4A). The breakdown of genomic feature categories in which peaks were located is presented in Table 2.

**Figure 4:**
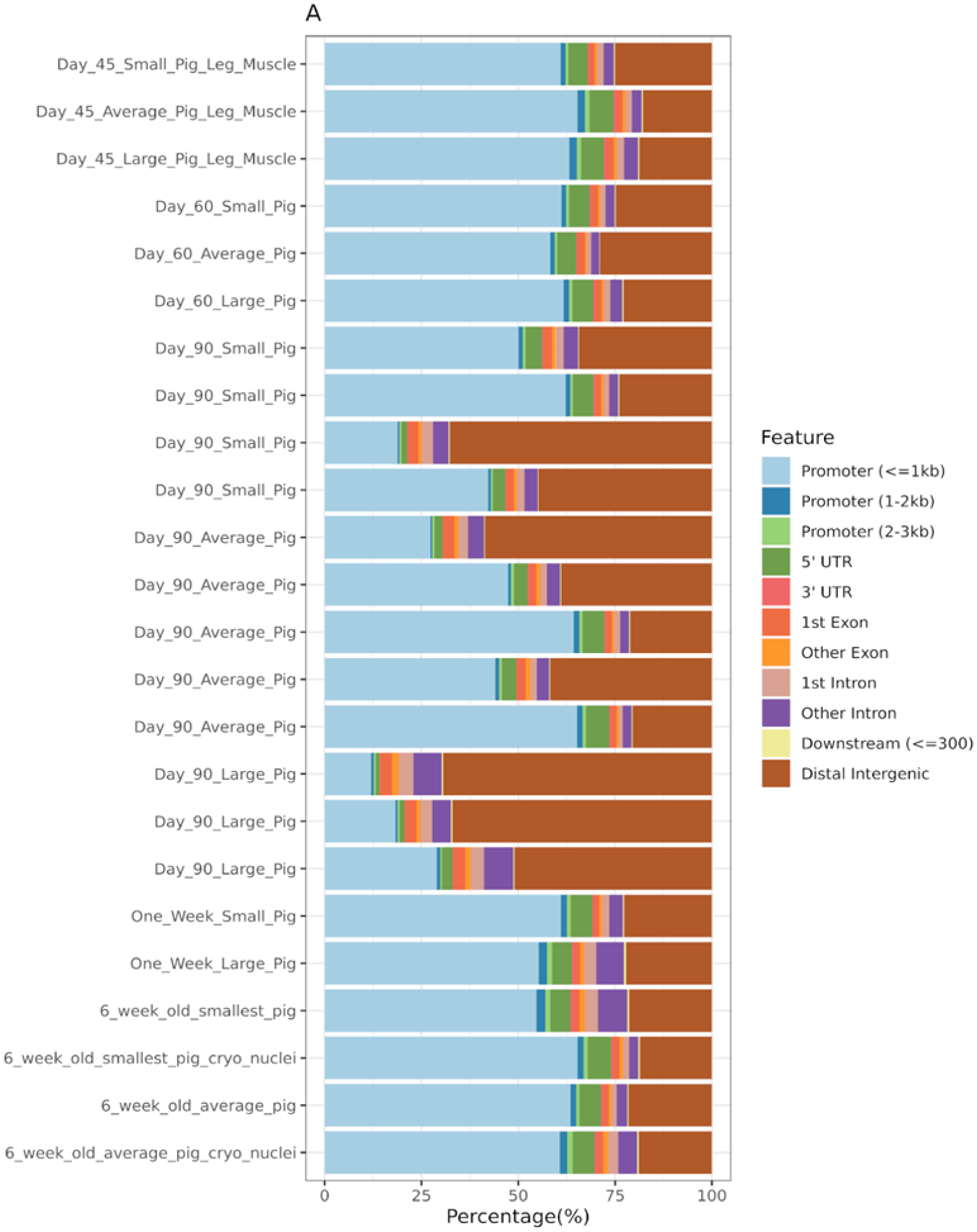
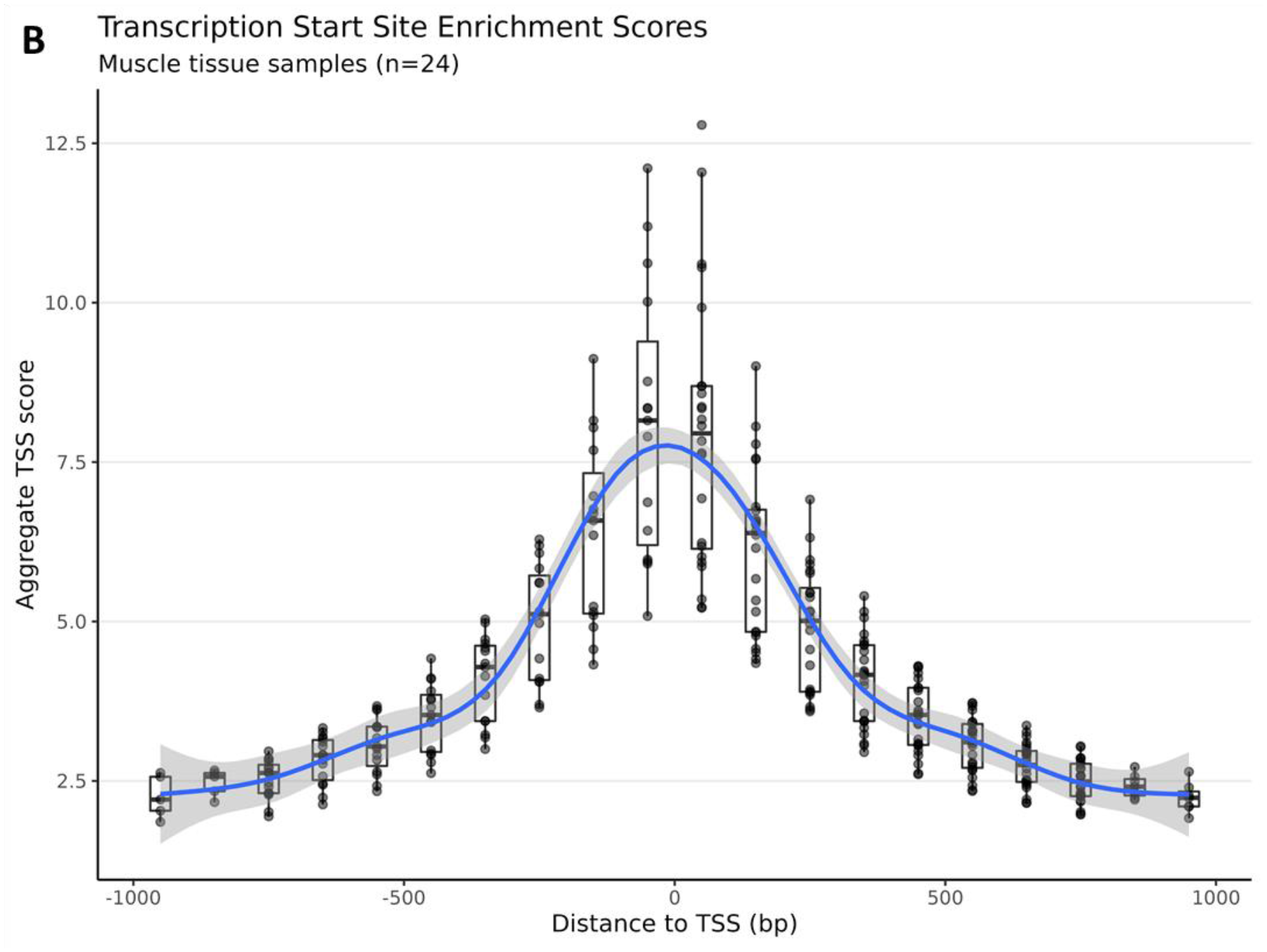
A) Percentage of ATAC-Seq peaks within genomic features. The samples are sorted by the developmental timeline (Day 45 to 6 weeks old from top to bottom. B) A boxplot graph of TSS enrichment score of ATAC-Seq peaks and their relative distance from the TSS for all 24 samples. TSS score = the depth of TSS (each 100bp window within 1000 bp flaking TSS) / the depth of flank ends (100bp each end). TSS-E score for each library = max(mean(TSS score in each window)) calculated by ATACseqQC v1.14.4 (Ou et al. 2018).

**Table 2.**
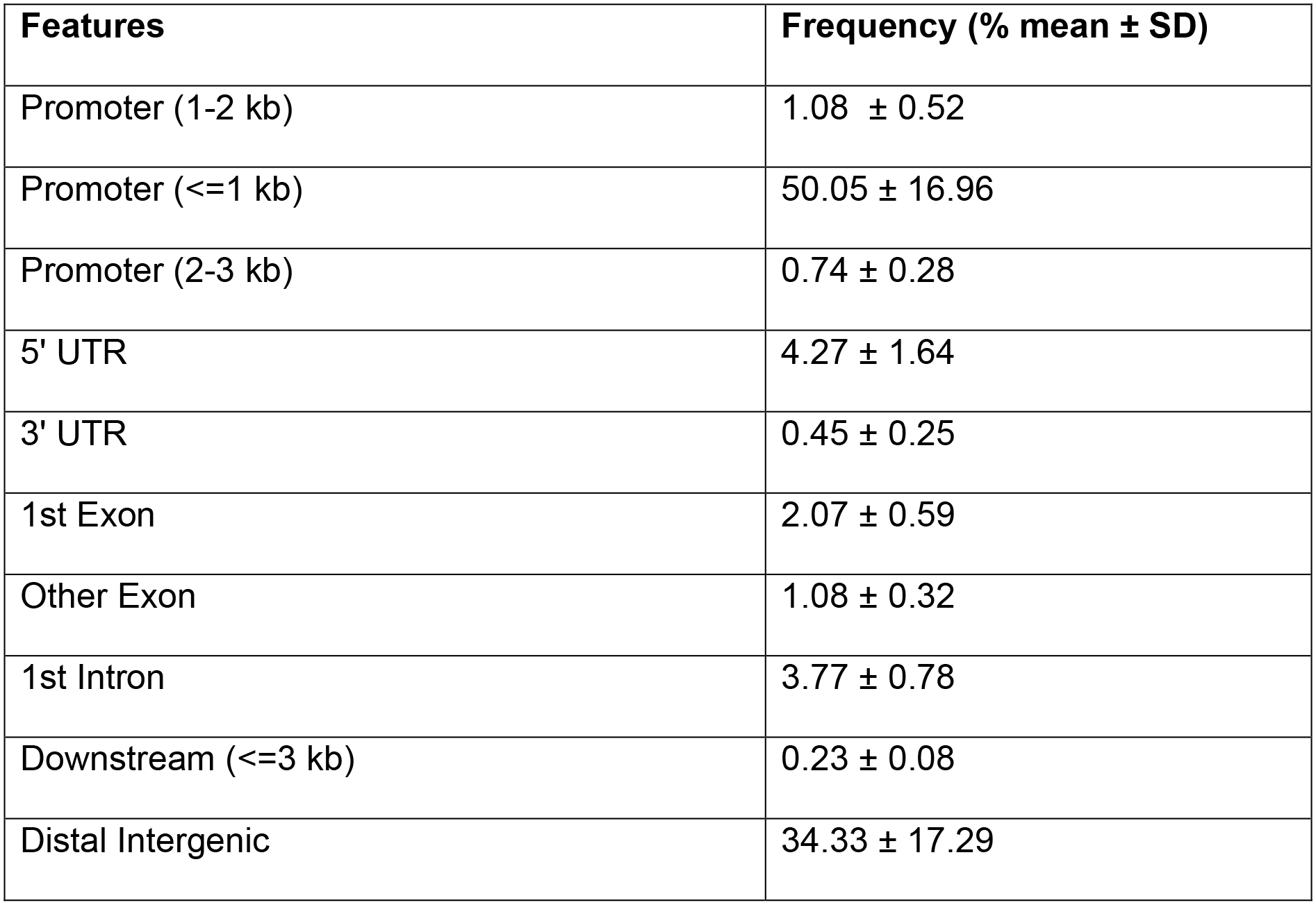
The frequency of ATAC-Seq peaks in each genomic feature category annotated by ChipSeeker and averaged across samples (3,766 annotated peaks from a total of 4,661 peaks).

### Proximity of ATAC-Seq peaks to transcription starts sites (TSSs)

The transcription start site enrichment score (TSS-ES) was used to validate the presence of the ATAC-Seq signal flanking TSS (Figure 4B). ATAC-Seq libraries had a TSS-ES of 8.03 ± 2.14 (mean ± SD) on average (min 5.21; max 12.8). As noted above there was a significant difference in the ATAC-Seq TSS-ES in the libraries from cryopreserved nuclei relative to the libraries from flash frozen tissues (the TSS-ES was higher in cryopreserved nuclei). All libraries showed a uniform distribution of the TSS scores flanking TSSs as shown in Figure 4B.

### Differential peak analysis of ATAC-Seq read counts using a consensus set of peaks

Differential peak analysis revealed 377 ATAC-Seq peaks from the consensus peak set in which the read count differed significantly between the developmental time points. These peaks were annotated using *Sscrofa11.1* corresponding to 724 unique transcripts (245 unique genes). 109 peaks were in unannotated intergenic regions. Nearly half of the peaks exhibiting differential read counts, between developmental time points, were located in intronic regions and only 11.1% resided in promoters as shown in Table 3.

**Table 3:** Genomic feature distribution of ATAC-Seq peaks where read counts were significantly different between the day 45, day 60, day 90, 1 week and 6 week time points.

| Features | Count | Frequency (%) |
| --- | --- | --- |
| Intronic | 185 | 49.1 |
| Intergenic | 109 | 28.9 |
| Promoter | 42 | 11.1 |
| 3' UTR | 12 | 3.18 |
| Proximal | 10 | 2.65 |
| 5' UTR | 8 | 2.12 |
| CDS | 8 | 2.12 |
| Exonic | 3 | 0.79 |

The read counts for the peaks that differed significantly between time points are shown in Figure 5 as normalised fragment counts. A detailed list of these peaks is included in Supplementary File 3. To test whether the fragment count distribution between the piglet sizes was different (Figure 5) we used ANOVA and found no significant differences between any of the pairwise comparisons (ANOVA + Tukey HSD, *p* value > 0.05).

**Figure 5:**
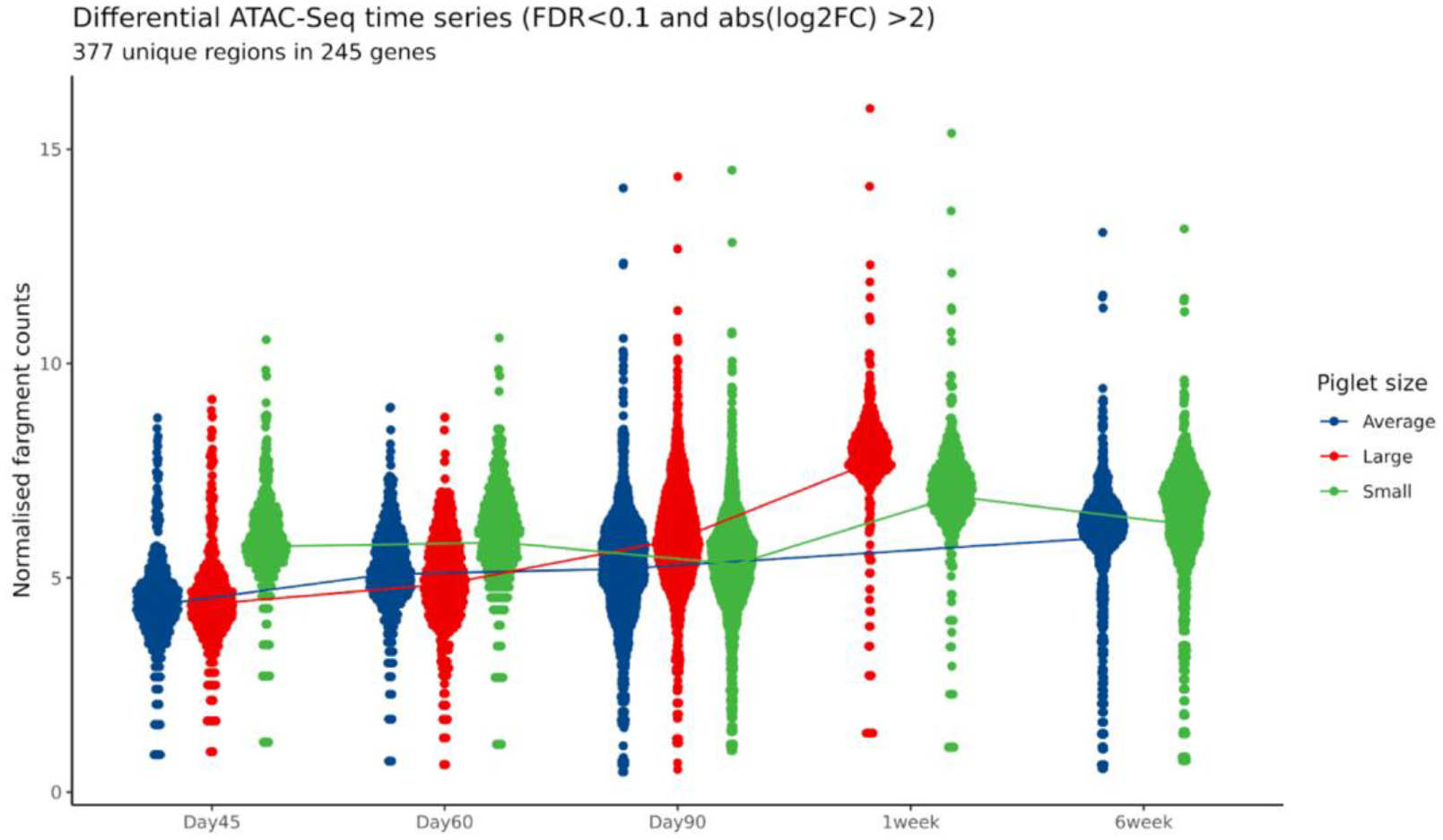
Normalised fragment counts plotted across developmental time points. A) Differential peak analysis of ATAC-Seq peaks across time points, including piglet size and litter as fixed variables in the DESeq2 LRT model. Significantly differentially expressed peaks (DEP) are plotted for each time point and coloured by piglet size. The line represents the average normalised read counts per time point for all DEPs. B) Shows the distribution of the DESeq2 normalised counts for the four ATAC-Seq peaks that were significantly differentially expressed between the piglet sizes at day 90, across the developmental time points.

### Transcription factor activity footprinting of the ATAC-Seq peaks (time and piglet size)

Transcription factor footprinting analysis across the developmental time points did not show any significantly different HINT scores (Figure 6A). In the comparison between large and small piglet size at day 90 samples, 5 differentially active transcription factors (TFs) (*GMEB2, TFAP2C(var.2), HOXD12, FOXH1* and *CEBPE*) were detected using JASPAR2020 database annotation. The TF *CEBPE* CCAAT-Enhancer-Binding Protein-Beta, showed the most extreme HINT z score (HINT z score-14.35) (Figure 6B). *CEBPE* is known to be upregulated after muscle injury and be highly associated with muscle strength in human and mouse models (Harries et al. 2012).However, the lack of visual evidence of a TF footprint in either small or large piglets (Figure 6B) indicates the extreme HINT z score might be the result of technical artefact. In comparison, *GMEB2,* a glucocorticoid receptor expression regulator (Kaul et al. 2000), was the only TF with significantly higher enrichment in the small size piglets (HINT z score 4.29) in comparison to the large piglets, and showed visual evidence of a TF footprint (Figure 6C). Details of the TF footprinting are shown in Table 4.

**Figure 6:**
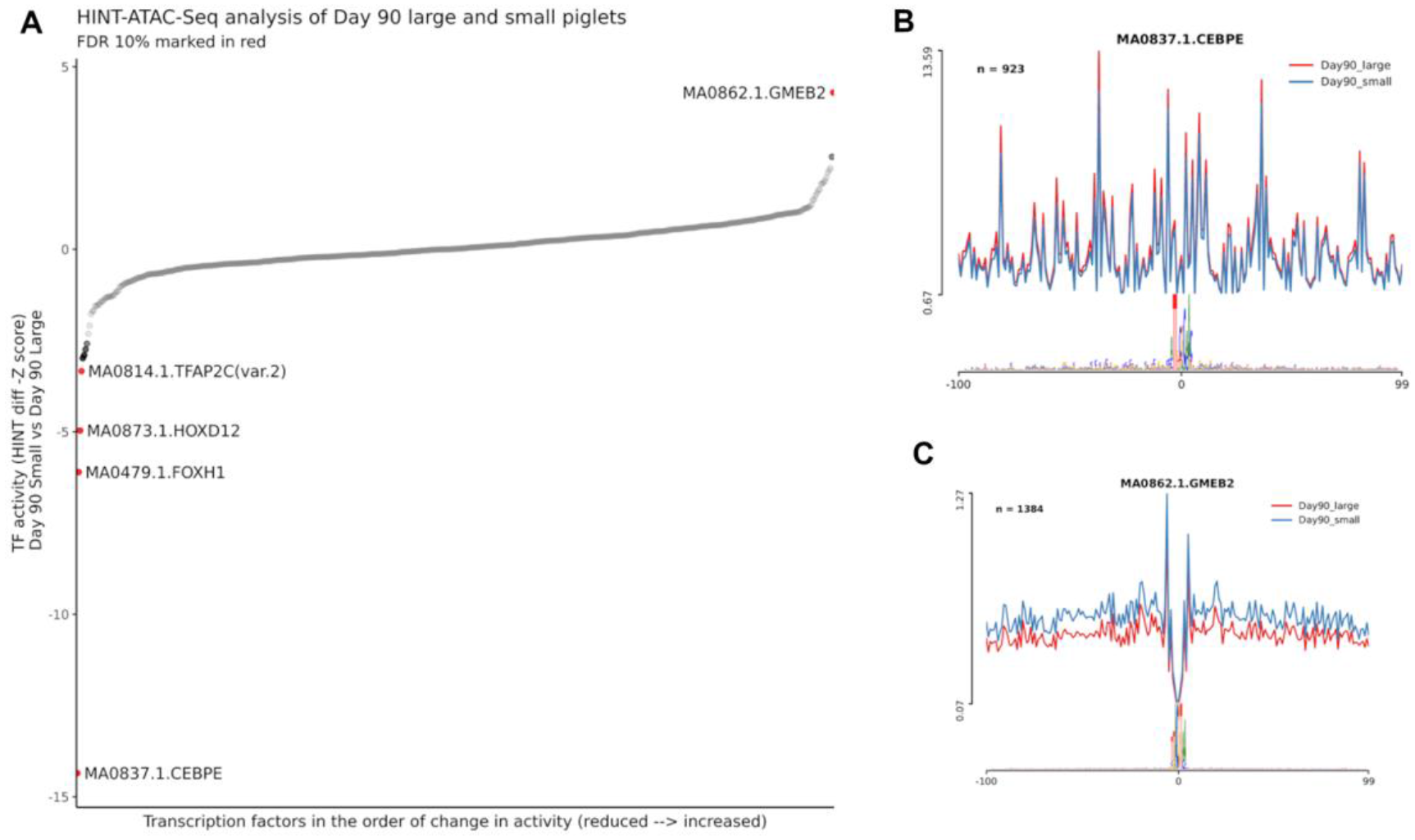
HINT pipeline analysis of the ATAC-Seq dataset for day 90 samples compared between large and small piglet size. A) Differential transcription factor activity between two piglet sizes at day 90 sorted by the HINT z-score value. The red dots are statistically significant (FDR 10%) showing hyperactivity of GMEB2 in the small size piglets muscle tissues along with lowered activity of TFAP2C, HOXD12, FOXH1 and CEBPE transcription factors (higher in the large size piglets). TF activity in the vicinity of the corresponding motif between large and small size piglets are shown in B) for CEBPE and C) for GMEB2.

**Table 4.** Transcription factor footprinting analysis of the ATAC-Seq dataset using HINT and the JASPAR annotation database. FDR: false discovery rate (10% was considered significant). Comparison was performed in day 90 samples Small to Large (S/L direction of activity value)

| **Motif** | **TF_Activity** | **Z_score** | **P_values** | **FDR** |
| --- | --- | --- | --- | --- |
| MA0837.1.CEBPE | -0.71 | -14.35 | 1.01E-46 | 5.59E-44 |
| MA0479.1.FOXH1 | -0.30 | -6.10 | 1.04E-09 | 2.87E-07 |
| MA0873.1.HOXD12 | -0.24 | -4.96 | 6.95E-07 | 1.28E-04 |
| MA0862.1.GMEB2 | 0.21 | 4.29 | 1.74E-05 | 2.4E-03 |
| MA0814.1.TFAP2C(var.2) | -0.16 | -3.33 | 8.49E-04 | 9.3E-02 |

### Analysis of gene expression using RNA-Seq

We generated RNA-Seq data from the same muscle tissue samples that were used to generate the ATAC-Seq libraries, in order to link regions of open chromatin with gene expression. The transcript expression estimates for the muscle tissue samples from the five developmental time points (26 samples in total) were calculated as Transcript per Million mapped reads (TPM) using Kallisto. The TPM expression estimates were then investigated using PCA (Figure 7) to identify any samples that did not group as expected according to developmental time point. The samples from each developmental time point clustered together as expected in the first two dimensions of the PCA (Figure 7).

**Figure 7:**
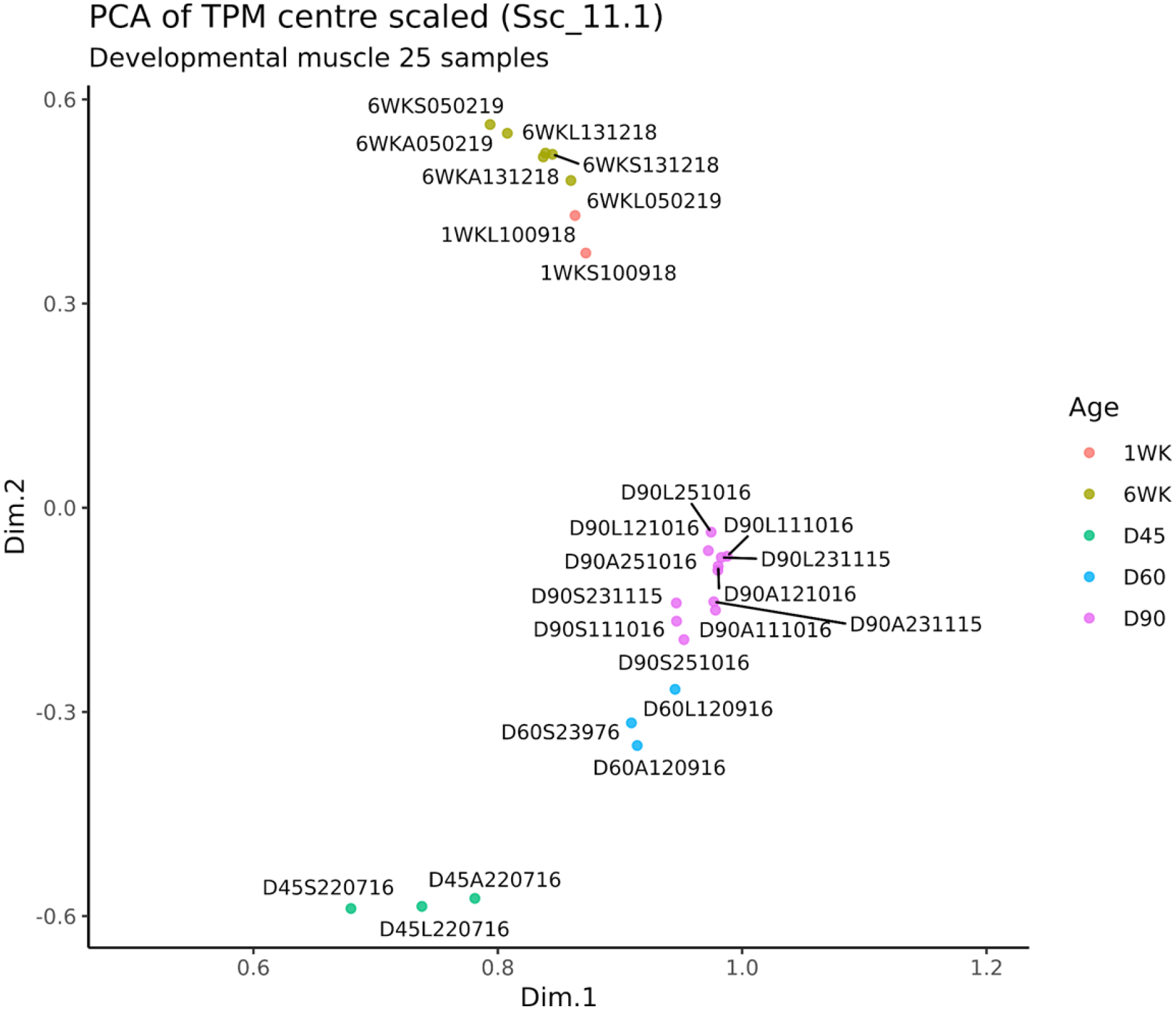
Principal Component Analysis (PCA) of gene expression estimates (as TPM) from the RNA-Seq data for each sample. The samples cluster according to developmental stage with clear separation of neonatal and post-natal samples. D45 = Gestational Day 45; D60 = Gestational Day 60; D90 = Gestational Day 90; 1WK = Neonatal 1 week old;6WK = Juvenile 6 weeks old.

### Analysis of genes that were differentially expressed between the three sizes of foetal piglet at day 90 of gestation

Differential gene expression analysis was performed using the TPM values for the three sizes of foetal piglet (small, average, large) at day 90 of gestation. Between the three sizes of foetal piglet 89 genes (FDR 10%), were found to be differentially expressed. When average vs small sized foetal piglets were compared, 58 up- and 31 down-regulated genes were detected. When large vs small sized foetal piglets were compared, 54 up- and 35 down-regulated genes were detected. Differentially expressed genes with an adjusted *p* value (FDR < 0.1) and log2 fold change (log2FC) >= 0.1 are annotated in Figure 8. The comparison between large and small sized foetal piglets resulted in the largest number (n=89) of differentially expressed genes. The list of differentially expressed genes and detailed analysis metrics can be found in Supplementary Table 5. Many of the genes that were differentially expressed between large and small and average and small foetal piglets are involved in skeletal muscle function and growth (Figure 8). The gene calsequestrin 1 (*CASQ1*), for example, which was 1.54 fold up-regulated (log2FC 0.63 ± 0.17 adjusted *p* value = 2.0e-02) in large relative to small foetal piglets is the skeletal muscle specific member of the calsequestrin protein family, and is highly expressed in skeletal muscle in adult pigs, see (http://biogps.org/pigatlas/) (Freeman et al. 2012; Summers et al. 2020). *MYBPC2*, a gene that encodes myosin binding protein C, was also 2 fold up-regulated (log2FC 1.02 ± 0.22 adjusted *p* value =3.78e-04) in large relative to small foetal piglets (Figure 8). It has also been shown to be highly expressed in the muscle of pigs, see (http://biogps.org/pigatlas/) (Freeman et al. 2012; Summers et al. 2020). The muscle specific transcription factor myogenin (*MYOG*) was down-regulated, (log2FC 0.28 ± 0.09 adjusted *p* value = 9.0e-02), in small relative to large foetal piglets (Figure 8).

**Figure 8:**
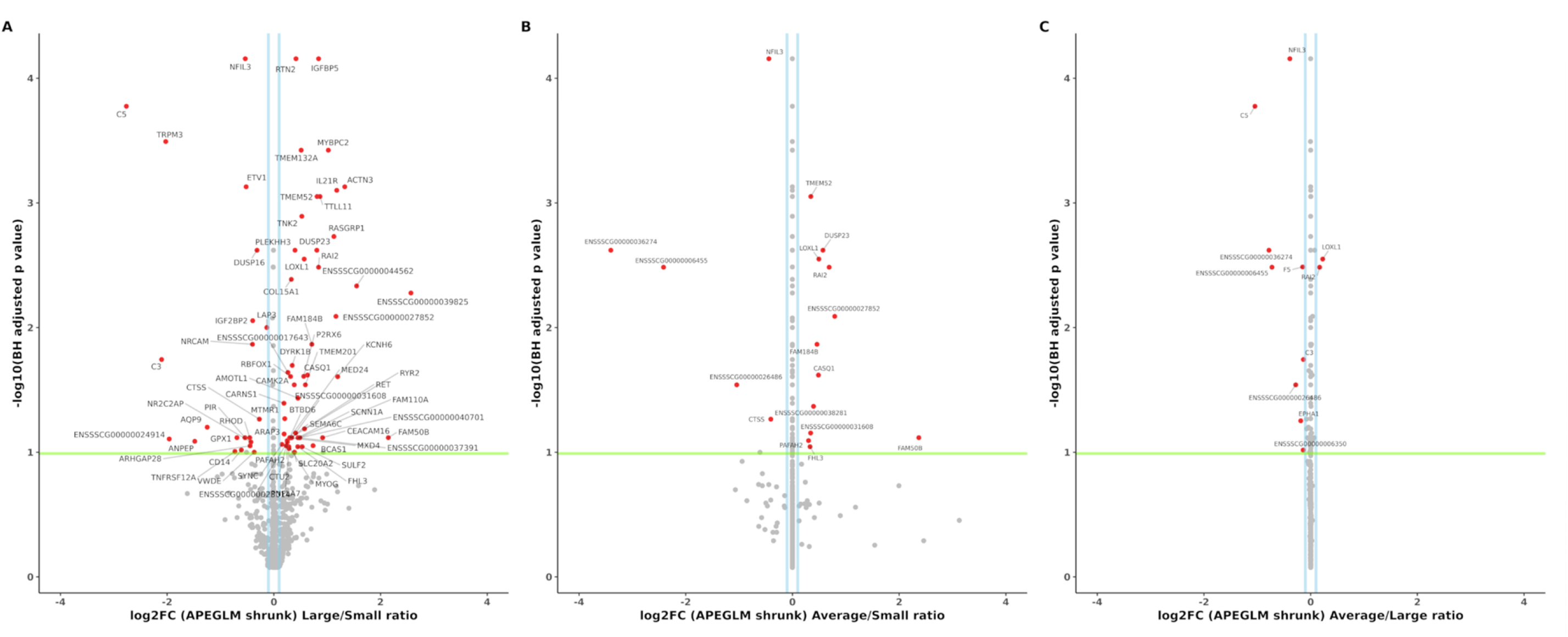
Differentially expressed genes (RNA-Seq) between large, average and small sized piglets at Day 90 of gestation. A) Large vs Small B) Average vs Small C) Average vs Large. Differentially expressed genes are shown in red, the log2FC >0.1 in blue and the significance threshold as a green line. The ApeGLM shrinkage method was used to normalised the log fold change plotted on the x axis as described in (Love et al. 2014).

### Overlay of the RNA-Seq differentially expressed genes and ATAC-Seq peaks from large vs small foetal piglets at day 90 of gestation

A further overlay of the ATAC-Seq and RNA-Seq datasets was performed for the day 90 large and small sized foetal piglets. ATAC-Seq peaks annotated using the Sscrofa11.1 Ensembl gene track information (black) and differentially expressed genes between the large vs small sized foetal piglets at day 90 (green) are shown in Figure 9. This analysis allowed us to determine which of the differentially expressed genes had an ATAC-Seq peak that was specific to either large or small sized piglets in its vicinity. The distribution of ATAC-Seq peaks around TSSs (within a 3 kb distance) was plotted for peaks specific to the large foetal piglets (Figure 9A), or specific to the small foetal piglets (Figure 9B). Size specific peaks within the 5’UTR region of four differentially expressed genes, *MYOG,* ryanodine receptor 2 (*RYR2*), transmembrane 4 L six family member 4 (*TM4SF4*) and interleukin 21 receptor (*IL21R*) (Figure 8), were only observed in the small foetal piglets (Figure 9B). There was no evidence of size specific peaks near these genes in the large sized foetal piglets (Figure 9B). Of the four genes, *MYOG* is known to be highly expressed in skeletal muscle tissue, see (http://biogps.org/pigatlas/) (Freeman et al. 2012; Summers et al. 2020). In some cases, a size-specific ATAC-Seq peak was located within the 5’ UTR region of a gene that was involved in muscle growth and down-regulated in small relative to large piglets. *MYOG*, for example, was down-regulated in small sized foetal piglets (Figure 8), with a regulatory region 315 bp in size 8,769 bp upstream of the TSS, that was present in the small sized piglets but absent in the large sized foetal piglets (Figure 9 A&B).

**Figure 9:**
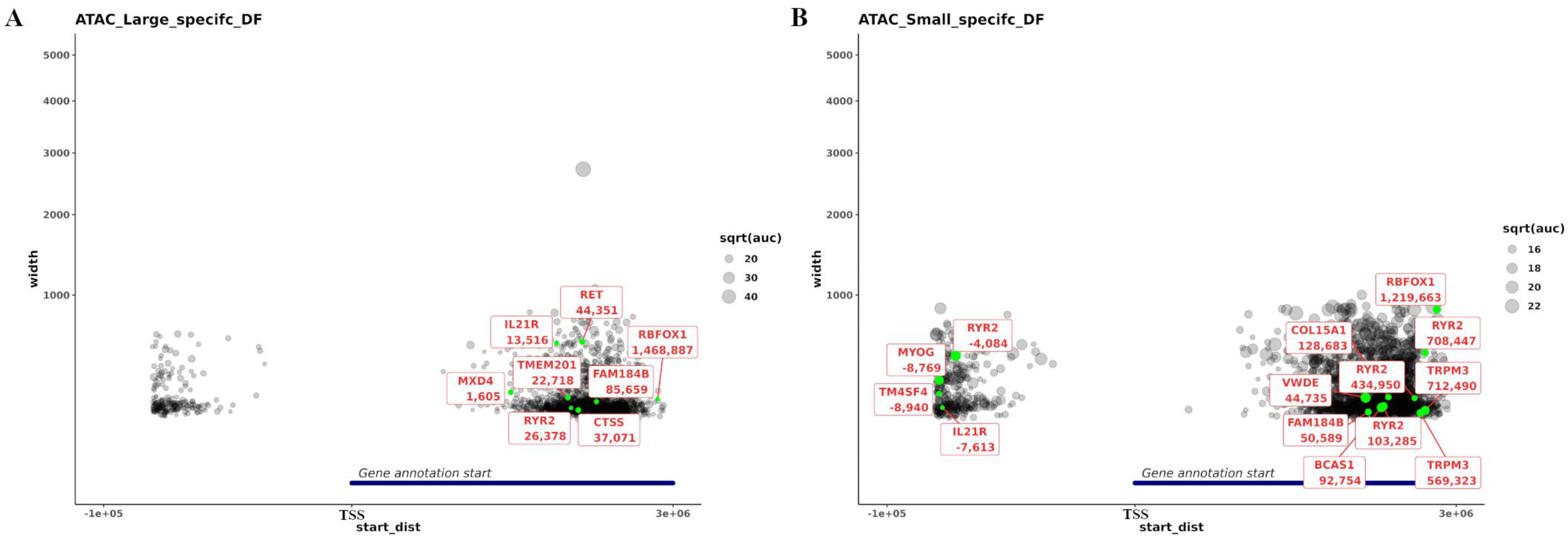
Proximity of ATAC-Seq peaks specific to large (A) and small (B) piglets and differentially expressed genes. Differentially expressed genes are marked in green. The x-axis (start_dist) is the distance from the start of the gene model to the start of the ATAC-Seq peak, for +ve values the peak is either within the gene or within 10 kb of the 3’ end and for -ve values the peak is within 10 kb of the 5’ end of the gene. The y-axis indicates the width of the peak. As the y axis represent the width of the peak, the larger the node the wider the ATAC-Seq peak.

## Discussion

In this study we used ATAC-Seq and RNA-Seq to improve our understanding of gene expression and regulation in developing pig muscle. The aims of the study were to: 1) Optimise the Omni-ATAC-Seq protocol for frozen pig muscle tissue; 2) Map regions of open chromatin in semitendinosus muscle tissue from small, average and large sized male piglets at five developmental stages (day 45, 60 & 90 of gestation, one and six weeks post-natal) and 3) Analyse RNA-Seq data from the same tissues to generate gene expression profiles.

To fulfil aim one, we optimised the Omni-ATAC-Seq protocol (Corces et al. 2017) for frozen muscle tissue. This, to our knowledge, is the first time the Omni-ATAC-Seq protocol (Corces et al. 2017) has been optimised for frozen tissue from a farmed animal species. Other studies have used ATAC-Seq to profile open chromatin in freshly sorted cell types e.g. (Foissac et al. 2019) or isolated and cryopreserved nuclei from dissociated tissue e.g. (Halstead et al. 2020a,b). Working with sorted cells was outside the scope of this study. We were, however, able to perform a comparison of ATAC-Seq libraries prepared from either flash frozen tissue or cryopreserved nuclei for a small subset of samples. We found that the datasets generated by the two methods were broadly comparable, in terms of the distribution of ATAC-Seq reads mapping to genomic regions. The libraries from cryopreserved nuclei had a higher TSS-Enrichment score in comparison to frozen tissues samples and this finding should be further validated with a larger number of samples.

The ability to utilise flash frozen tissue effectively for ATAC-Seq is advantageous for two reasons. Firstly, many legacy samples from large animal studies have been flash frozen then archived at −80°C and represent a very valuable resource if they can be utilised. Secondly, flash freezing is straightforward to undertake and standardise, especially when the logistics of collecting samples from large animals can be technically challenging (Wong et al. 2012). For this study we have only optimised the Omni-ATAC-Seq protocol for flash frozen muscle tissue samples from pigs. Expanding the protocol to other tissues and other species should be relatively straightforward, although some tissue-specific optimisation will be required, particularly for tissues that are known to be complex to work with.

For aim two of the study, we generated open chromatin profiles, in the form of ATAC-Seq peaks, for *semitendinosus* muscle from piglets from five developmental stages. The developmental stages were chosen according to their relevance to muscle development. ATAC-Seq peaks mapped as expected to promotor regions and within 1 kb of the TSS, which is consistent with studies across different species (Foissac et al. 2019; Yue et al. 2021). A study of *longissimus dorsi* muscle from pig embryos at days 45, 70 and 100 conducted by (Yue et al. 2021) showed that 30%, 21%, and 14% of the peaks were identified in promoter regions respectively. Of these peaks, 91% mapped to within −1 kb and +100 bp of the TSS (Yue et al. 2021). A cross-species analysis of ATAC-Seq data showed that in mice, goats, cattle, pigs and chicken, 10-15% of ATAC-Seq peaks were located within up to 5 kb of the TSS, and were therefore considered as promoters (Foissac et al. 2019). The results from our study showed that a majority of the ATAC-Seq peak frequency was located within ±1 kb of the TSS, with the remaining primarily located within distal intergenic regions.

The distribution of ATAC-Seq peaks in intergenic regions at day 90 in large and small sized piglets indicated that piglets of different sizes show changes in genome regulation primarily at intergenic sites (i.e. differential enhancer activity). Day 90 is a critical stage of muscle development when fibre formation ceases and muscle growth accelerates through fibre hypertrophy (Oksbjerg et al. 2004). Significant up-regulation of genes involved in muscle growth also occur at day 90 (Zhao et al. 2015; Ayuso et al. 2016; Yue et al. 2021). As such chromatin may be more open at this developmental stage to allow transcription factor binding prior to the rapid muscle growth that occurs during the early postnatal period (Rudar et al. 2019). (Yue et al. 2021) reported widespread increases in accessible chromatin and increasing regulatory complexity in developing pig embryos through days 45, 75 and 100 of gestation. Studies profiling open chromatin in preimplantation embryos found global differences in chromatin accessibility between embryo stages in humans (Wu et al. 2018; Liu et al. 2019) and cattle (Halstead et al. 2020c). ATAC-Seq datasets for post implantation embryos in humans and other mammalian species are limited. ChIP-seq analyses of a wide variety of histone markers in the brain, heart, and liver of early human embryos identified developmental stage-specific patterns in the epigenome (Yan et al. 2016). The sample size in our study was small, with only a few biological replicates for most points, with the exception of Day 90. Even so, our results, are in agreement with other recent studies e.g. (Yue et al. 2021), and indicate that chromatin accessibility and regulation of gene expression changes throughout development in the pig muscle. This is significant for studies aiming to understand when during development functional variation in the genome has an effect on the adult phenotype. For example, in this study, transcription factor footprint analysis, showed that the transcription factor *GMEB2,* which increases sensitivity to glucocorticoids (Kaul et al. 2000), had significantly higher TF activity in small relative to large size piglets at day 90 of gestation. This finding is potentially phenotypically relevant because low birth weight piglets have been shown to have higher in utero-cortisol levels than their normal birth weight litter mates (Roelofs et al. 2019).

To address aim three, we analysed gene expression information for the same muscle tissue samples. We used this approach to compare gene expression and chromatin openness between foetal piglets of different size at day 90 of gestation. Other studies have used a similar approach to investigate the effect of histone modification on the expression of genes involved in placental development in pigs (Han et al. 2019) and chromatin accessibility in pre-natal muscle development (Yue et al. 2021). Differences in open chromatin were reflected in the expression of genes involved in muscle growth. Analysis of the RNA-Seq data revealed that genes associated with muscle growth, including *CASQ1, MYBPC2* and *MYOG*, were differentially expressed in large relative to small piglets. Differential expression of myogenic genes (*e.g. MYOG*) in pig muscle has been previously reported by (Felicioni et al. 2020) who compared intrauterine growth restricted and normal weight piglets. *CASQ1* encodes the skeletal muscle specific member of the calsequestrin protein family, is related to muscle metabolism, and has been shown to be highly expressed in fat pig breeds (Zhao et al. 2011). In this study, *CASQ1* was up-regulated in large relative to small foetal piglets. *MYBPC2* encodes the fast isoform of the myosin binding protein C family (Weber et al. 1993). In Piedmontese (*GDF8* mutant) cattle *MYBPC2* is highly expressed in foetal muscle, reflecting fast glycolytic fibre structural differentiation (Lehnert et al. 2007). In this study, *MYBPC2* was highly up-regulated in large versus small foetal piglets, potentially reflecting a greater proportion of fast glycolytic muscle fibres.

*MYOG* was down regulated in small sized foetal piglets relative to large sized foetal piglets. *MYOG* is essential for myoblast fusion during muscle development (https://www.uniprot.org/uniprot/P49812) and associated with QTLs, for body weight at birth (https://www.animalgenome.org/cgi-bin/QTLdb/SS/qdetails?QTL_ID=8656) and backfat thickness (https://www.animalgenome.org/cgi-bin/QTLdb/SS/qdetails?QTL_ID=8657), according to Genome Wide Association Studies (GWAS) (Xue and Zhou 2006). Other studies measuring gene expression also found *MYOG* was down-regulated in muscle cell types from low birth weight piglets (Felicioni et al. 2020; Stange et al. 2020) and in pigs with high levels of intramuscular fat (Lim et al. 2017). When we compared the ATAC-Seq and RNA-Seq data we identified an ATAC-Seq peak within the 5’ UTR region of *MYOG* (315 bp in size, 8,769 bp upstream of the TSS) that was present in the small foetal piglets but missing from the large foetal piglets for Day 90 of gestation. In future work we plan to remove this peak using CRISPR genome editing and measure the effect on primary muscle cells in culture. Further validation of the variants within this regulatory region would be useful to determine whether they might underlie variation in intramuscular fat or myofiber specification as was the case for the regulatory variant recently characterised in *MYH3* (Cho et al. 2019). Open chromatin regions in the vicinity of genes that were significantly differentially expressed in large relative to small piglets were of interest in this study, and other similar studies e.g. (Yue et al. 2021) particularly because they are located near TSS or promoter coordinates. Further functional validation is required to determine whether chromatin accessibility has any direct effect on the expression level of differentially expressed genes in this context.

The datasets we have generated for this study provide a foundation for incorporating functional information in statistical analyses, to increase the precision and power with which we can fine map high quality causal variants in pigs. This would make it possible to increase the accuracy of genomic selection and the efficiency with which breeding turns genetic variation into genetic gain. The next stage of the study is to leverage the ATAC-Seq and RNA-Seq data with a very large dataset of genetic variants from production pigs to determine whether any trait-linked variants are located within the open chromatin regions we have identified for muscle tissue. The characterisation of regulatory and expressed regions of the genome in muscle tissues also provides a basis for genome editing to promote functional genomic variants in pig breeding programmes (Jenko et al. 2015; Hickey et al. 2016; Johnsson et al. 2019), providing a route to application for FAANG data.

## Conclusions

The dataset we have generated provides a powerful foundation to investigate how the genome is regulated in production pigs and contributes valuable functional annotation information to define and predict the effects of genetic variants in pig breeding programmes. The outcomes of the study will: 1) help us to understand the molecular drivers of muscle growth in pigs; 2) provide a foundation for functionally validating target genomic regions in *in vitro* systems and 3) identify high quality causative variants for muscle mass with the goal of harnessing genetic variation and turning it into sustainable genetic gain in pig breeding programmes.

## Acknowledgements

The authors would like to thank the staff and farm technicians at both Dryden Farm and Easter Howgate. Charis Hogg provided assistance in planning and collection of samples from the foetal time points and Christopher Proudfoot coordinated sampling of the six-week-old pigs. ATAC-Seq and RNA-Seq libraries were sequenced by Edinburgh Genomics. The authors are also grateful for the support of the FAANG Data Coordination Centre (http://data.faang.org) in the upload and archiving of the sample data and metadata.

## Ethics Statement

All projects from which samples for this study were collected were reviewed and approved by The Roslin Institute, University of Edinburgh’s Animal Work and Ethics Review Board (AWERB). All animal work was carried out under the regulations of the Animals (Scientific Procedures) Act 1986.

## Data Availability

The raw sequence data for the ATAC-Seq samples (n=24) are available via the European Nucleotide Archive (ENA) under accession number PRJEB41485. Details of all samples processed for the RNA-Seq dataset (n=26) can be accessed via the ENA under accession number PRJEB41488. The sample metadata is available via the BioSamples database under sample accession numbers SAMEA7178119, SAMEA7178120, SAMEA7178122, SAMEA7178123, SAMEA7178124, SAMEA7178125, SAMEA7178126, SAMEA7178127, SAMEA7178134, SAMEA7178138, SAMEA7178142, SAMEA7178149, SAMEA7178150, SAMEA7178153, SAMEA7178159, SAMEA7178160, SAMEA7178164, SAMEA7178178, SAMEA7178179, SAMEA7178180, SAMEA7178182, SAMEA7178183, SAMEA7178184, SAMEA7178185, SAMEA7178187 and SAMEA7178188. These datasets are curated and submitted to FAANG data portal according to FAANG’s sample and experimental guidelines (Harrison et al. 2018). All the sample, experiment and analysis protocols for this study are also available through the FAANG Data Coordination Centre via the following links: https://data.faang.org/api/fire_api/samples/ROSLIN_SOP_ATAC_Seq_DNAIsolationandTagmentation_Frozen_Muscle_Tissue_20200720.pdf, https://data.faang.org/api/fire_api/samples/ROSLIN_SOP_ATAC-Seq_DNAIsolationandTagmentation_Cryopreserved_Muscle_Nuclei_Preparations_20200720.pdf, https://data.faang.org/api/fire_api/samples/ROSLIN_SOP_ATAC-Seq_LibraryPreparationandSizeSelection_20200720.pdf, https://data.faang.org/api/fire_api/samples/ROSLIN_SOP_RNA_IsolationoftotalRNAfromfrozentissuesamples_20200720.pdf, https://data.faang.org/api/fire_api/experiments/ROSLIN_SOP_ATAC-Seq_Cryopreservednucleisamplesfrompigmuscletissue_Experimental_Protocol_20200720.pdf, https://data.faang.org/api/fire_api/experiments/ROSLIN_SOP_ATAC-Seq_Frozenpigmuscletissuesamples_Experimental_Protocol_20200720.pdf, https://data.faang.org/api/fire_api/experiments/ROSLIN_SOP_RNA_RNAIsolationandSequencingofPigMuscleTissues_Experimental_Protocol_20200720.pdf, https://data.faang.org/api/fire_api/analyses/ROSLIN_SOP_ATAC-Seq_analysis_pipeline_20201113.pdf, https://data.faang.org/api/fire_api/analyses/ROSLIN_SOP_RNA-Seq_analysis_pipeline_20201113.pdf, https://data.faang.org/api/fire_api/samples/ROSLIN_SOP_Collection_of_tissue_samples_for_ATAC-Seq_and_RNA-Seq_from_large_animals_20200618.pdf, https://data.faang.org/api/fire_api/samples/ROSLIN_SOP_Cryopreservation_of_Nuclei_for_ATACSeq_using_GentleMACS_20201119.pdf. All the supplementary Tables, Figures and files are also available from https://doi.org/10.6084/m9.figshare.13562285.

## Author Contributions

ELC devised the study and acquired the funding with MAH, FXD and ALA. CJA and CS designed the experiment from which the foetal tissues were collected. YCA, FXD, CS and CJA collected the samples from the foetal and one week old piglets. MS and ELC collected the samples from the six week old piglets. MS and ELC performed the cryopreservation of nuclei. ELC and MAH performed the ATAC-Seq library preparation. ELC and YCA extracted the RNA. MMH provided advice on ATAC-Seq library preparation and performed the transcription factor footprinting analysis. SAW collated all sample metadata for the project. MS performed all bioinformatic and data analysis. ELC wrote the manuscript with MS. MJ provided critical assessment of the manuscript. All authors read and approved the final version.

## Code Availability

The bioinformatic pipelines used for processing the ATAC-Seq (mapping and peak calling), RNA-Seq (transcript level expression analysis) and Allele specific expression are available via a code repository at https://msalavat@bitbucket.org/msalavat/pig_muscle.git respectively (https://bitbucket.org/msalavat/pig_muscle/src/master/).

## Supplemental Material

Supplementary Table 1: Weights and crown-rump lengths of all piglets included in the study.

Supplementary Table 2: Details of components of all buffers used for ATAC-Seq sample collection and library preparation.

Supplementary Table 3: Details of the primers (Ad2.x variable index) used for generating each ATAC-Seq library. Each sample is barcoded with a different variable index.

Supplementary Table 4: Summary table of RNA-Sequencing and ATAC-Sequencing quality control metrics for all samples. ‘N/A’ indicates that sequencing did not occur for that sample.

Supplementary Table 5: Results of differential expression analysis of large vs average vs small piglet sizes at Day 90 of gestation using a DESeq2 glm model. Only genes with FDR < 0.1 have been included.

Supplementary Table 6: QC metrics recommended by the ENCODE for all 24 ATAC-Seq libraries.

Supplementary_file_S1.zip: The collection of scripts and BED files corresponding to the ATAC-Seq peak coordinates for the sizes of piglets at Day 90.

Supplementary_file_S2.zip: The library preparation metadata and sequencing read depth metrics of the ATAC-Seq samples (n=24).

Supplementary_file_S3.zip: The differential peak analysis outputs for timeseries and Day 90 piglet size comparisons.

Supplementary Figure S1: Average genomic coverage of the ATAC-Seq dataset by chromosome.

Supplementary Figure S2: Genomic track visualisation of the ATAC-Seq dataset by raw read coverage at the gene coordinates for *GAPDH*.

Supplementary Figure S3: Metrics to compare ATAC-Seq peaks in libraries prepared from either cryopreserved nuclei from fresh tissue (blue) or from flash frozen tissue (red). Both sets of samples were collected from the semitendinosus muscle from the same six-week-old piglets. The results of a t.test comparison of the protocols for each of the genomic feature categories is presented in the enclosed table.

Supplementary Figure S4: Scatter plots of the Nucleosome free region score (NFR-score) flanking TSSs in all ATAC-Seq libraries (n=24). The Sus scrofa 11.1 transcript models were used for calculating NFR score flanking TSSs in ATACseqQC v1.14.4 package in R. The nucleosome free (middle 100bp of the TSS window) score of 0 is coloured differently from regions with nf score > 0.

