## Supplementary figures and images for "Profiling of open chromatin in developing pig (*Sus scrofa*) muscle to identify regulatory regions"

### Supp Fig 1

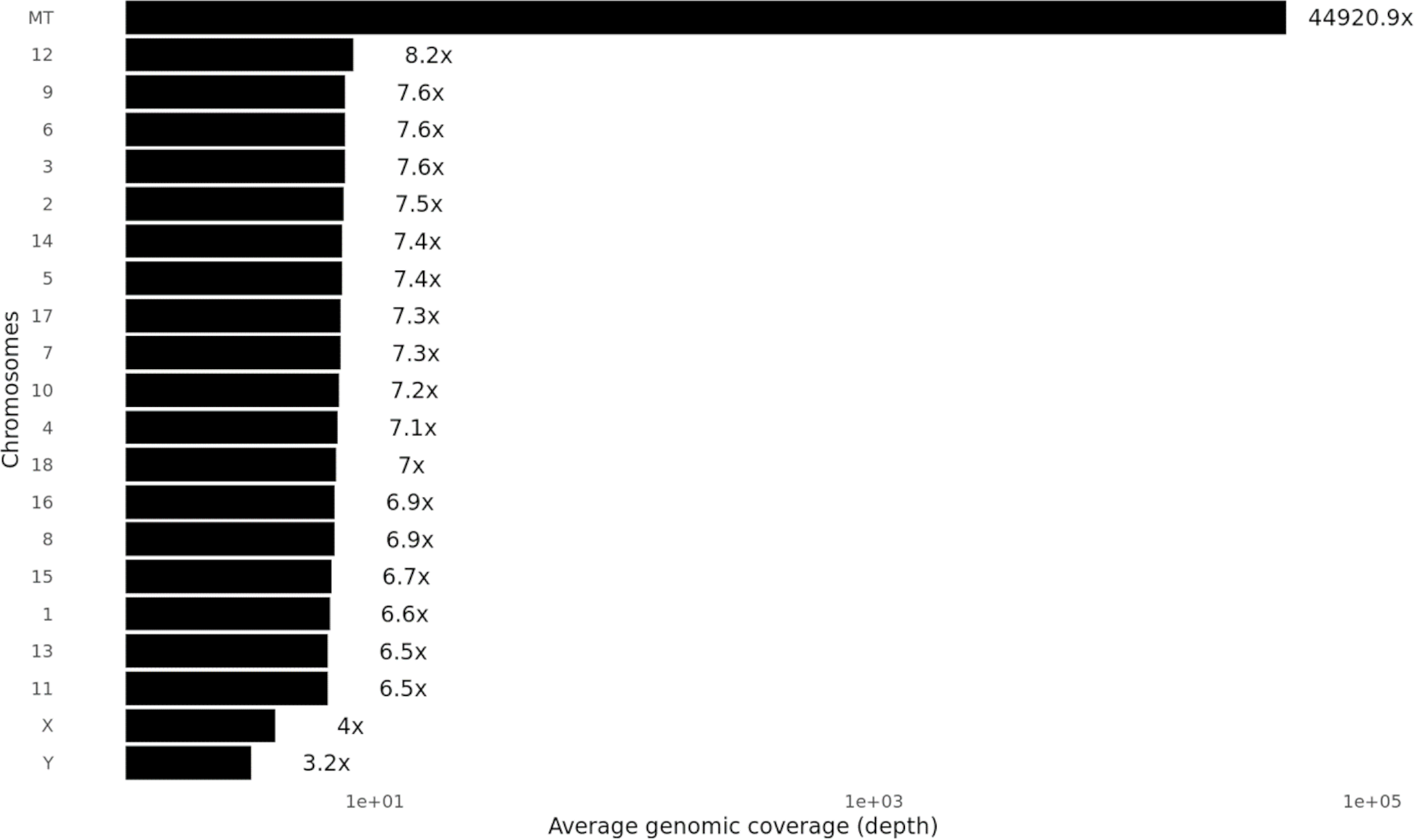

### Supp Fig 2

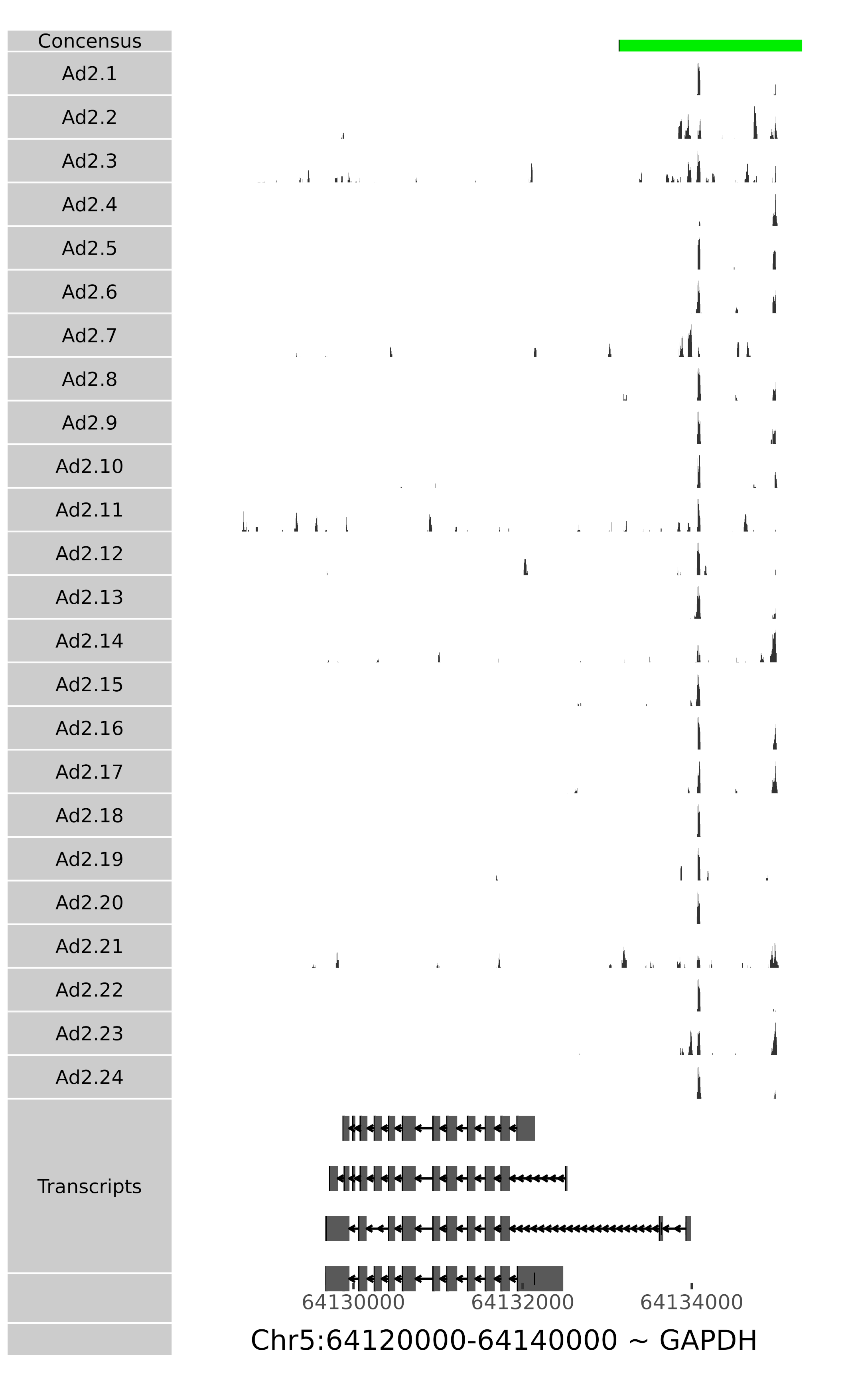

### Supp Fig 3

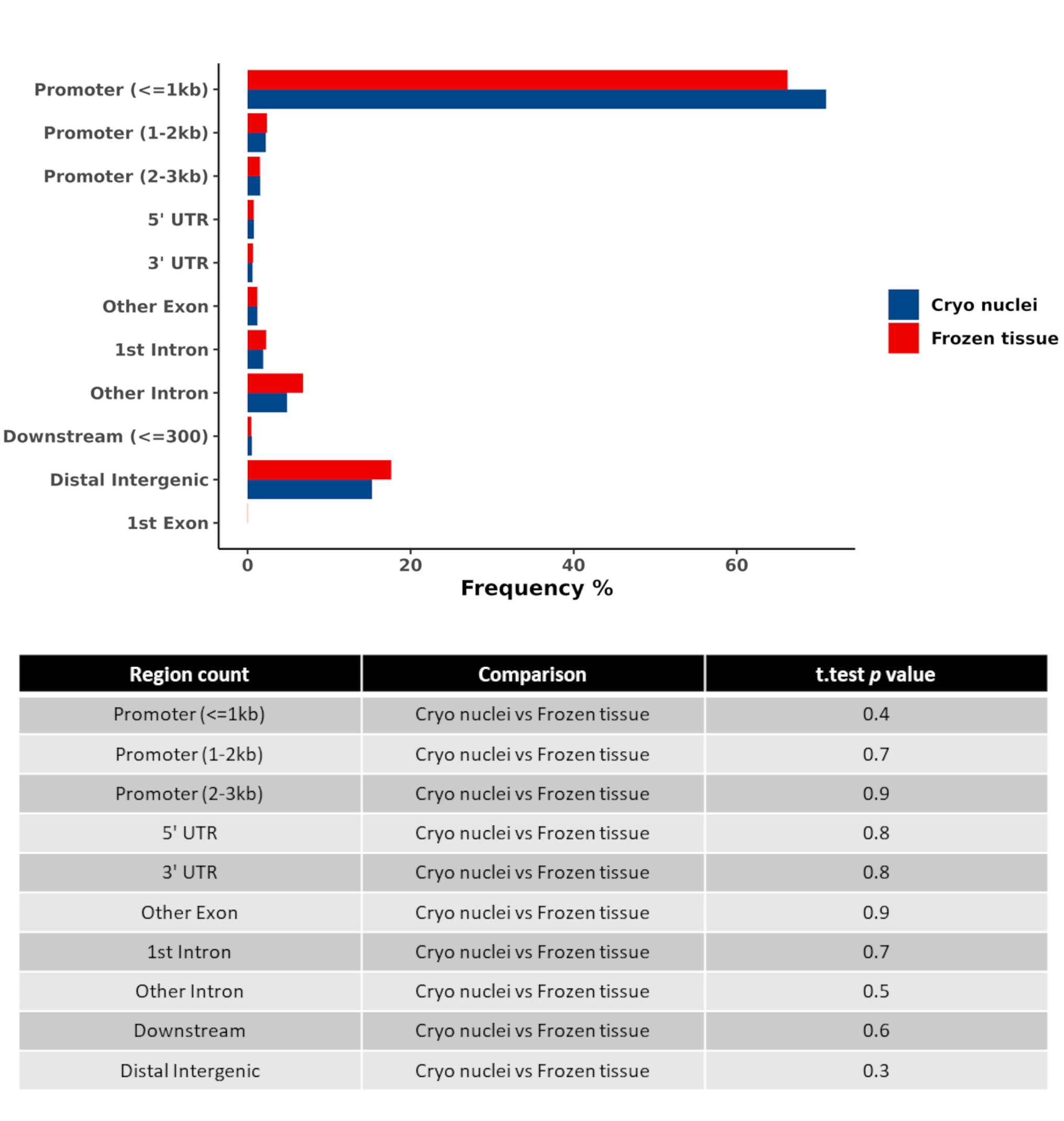

### Supp Fig 4

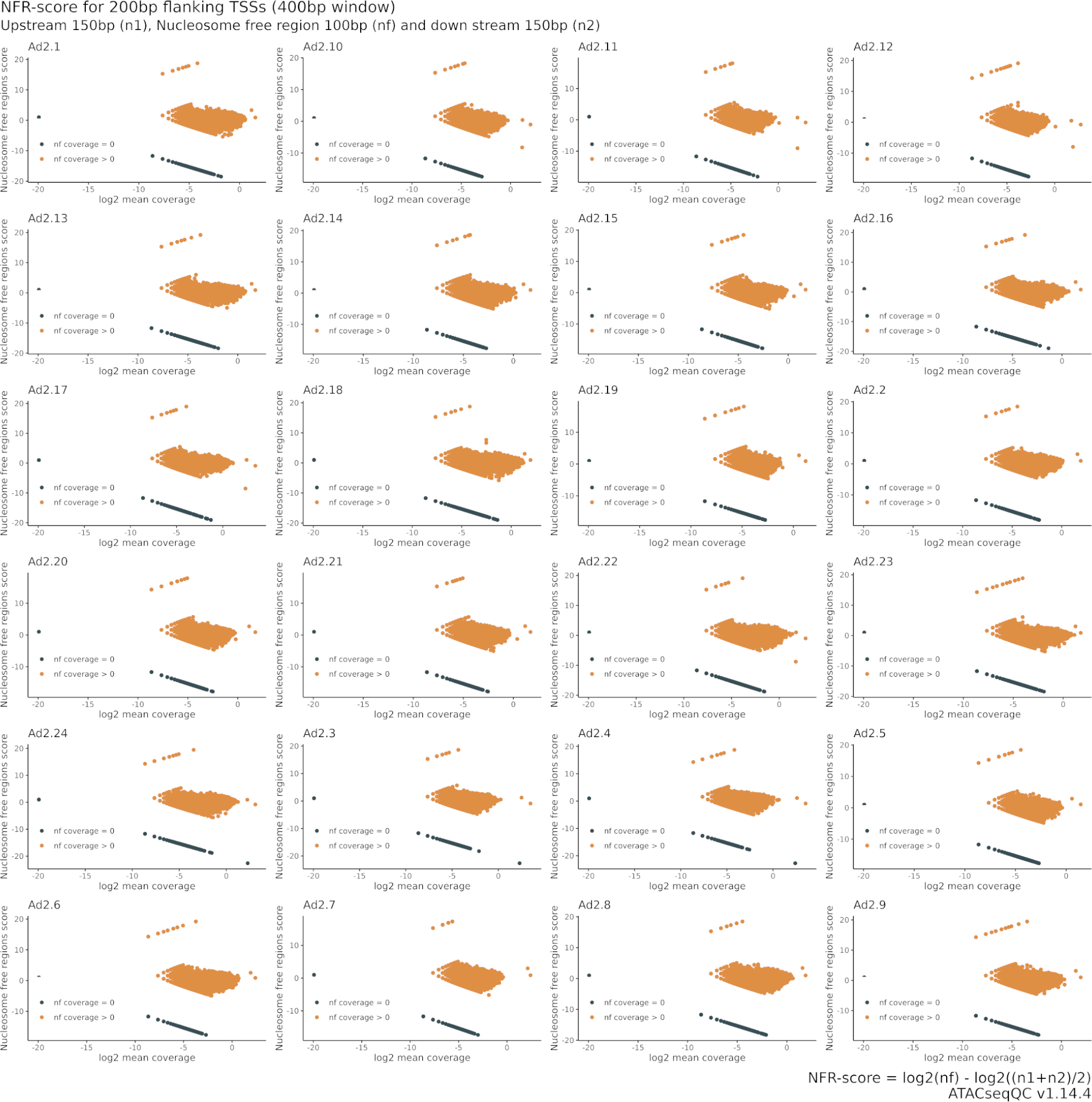
